# The Natural Material Evolution and Stage-wise Assembly of Silk Along the Silk Gland

**DOI:** 10.1101/2024.04.16.589504

**Authors:** Ori Brookstein, Eyal Shimoni, Dror Eliaz, Nili Dezorella, Idan Biran, Katya Rechav, Ehud Sivan, Anna Kozell, Ulyana Shimanovich

## Abstract

Silk fibers, with their highly ordered structure and mechanically superb properties, are produced in arthropod glands at minimal energy input and ambient conditions, a remarkable feat yet to be achieved synthetically. Due to the high instability and shear sensitivity of the silk protein feedstock, understanding silk fiber formation has been largely limited to *in-vitro* studies of certain gland sections, offering only a fragmented view of this process. Here, we monitor the whole silk feedstock processing *in-situ*, at the nano- to micron-scales, through imaging its progressive macromolecular assemblies and phase transitions along the entire *Bombyx mori* silkworm silk gland. This is done by combining state-of-the-art microscopy techniques, such as cryogenic sample preparation, fixation, and imaging. Our work reveals that fibroin assembles into micron-sized spherical storage “compartments” in the posterior and middle gland sections, a state that ensures its stability and avoids premature fibrillation. These compartments undergo several structural transformations along the gland and eventually disassemble at the entry to the anterior section, before the silk feedstock spinning begins. The spinning itself commences via a series of structural transitions, from the alignment of protein chains in liquid feedstock, through the formation of several fibrillated nano-structures and, in the final stage, a network of cross-linked nano-bundles, which determines the structure and properties of the final microfiber. Importantly, the length of the anterior section of the silk gland enables such gradual and balanced structural transitions. This direct imaging of silk’s natural formation process can help formulate a template for the transformation of fibrillar proteins into synthetic bio-fibers.

**Dedication:** This work is dedicated to the memory of Dr. Eyal Shimoni, who was a valued colleague and a dear friend. Eyal was a vital part of this research and was essential in shaping its direction. He will be deeply missed for his intellect, mindfulness, creativity, and unwavering dedication to scientific development. Though he is no longer with us, his influence and spirit continue to inspire us in our scientific pursuits. May his passion for discovery and commitment to excellence live on through this work.

## Introduction

Many natural materials exhibit exceptional properties and unique functionalities that have yet to be mimicked by industry. Silk is one such biomaterial. It is a mechanically superb, hierarchically ordered protein-based fiber produced by the silk gland of many organisms, including silkworms, spiders, and aquatic insects ^1–3^. Emulating the physical performance and production process of silk fibers is a great aspiration of biomimetics. Yet, thus far, only a partial understanding of the silk fiber formation process has been achieved.

The exceptional mechanical characteristics of silk fibers, which include a high Young’s modulus, strength, and toughness, originate from their hierarchically ordered structure^4–9^, which can vary widely between silk-spinning organisms^10–12^. Among those, the dragline spider silk fibers stand out for their superb mechanical performance. Despite differences in physiology among silk-spinning organisms (for example, spiders can have seven different silk glands, while silkworms have two), there are many similarities between their silk glands. They all consist of a protein-synthesis (posterior) section, a large feedstock storage (middle) section, and a long, narrow spinning (anterior) section^13–19^. The overall processing of silk feedstock is also usually quite similar, beginning with an unordered liquid protein that transitions into a β-sheet-rich solid fiber^6,15,20,21^. Some of the main structural differences between silk fibers of different silk-spinning species lie in the size and shape of the nano-fibrils, the way they are organized in the fiber, and the molecular structure dictated by the composing silk protein, unique to each organism^10–12^. The exact hierarchy of silk fibers is an ongoing debate and includes diverse structural models^2,22–25^, however, it is overall agreed that most types of silk fibers are bundles composed of aligned protein nano-fibrils^10,11,26^.

Silk of the domesticated *Bombyx mori* (*B. mori*) silkworms, which is the focus of our current investigation, are spun as a bave of two fibroin fibers coated by a layer of glue-like glycoproteins called sericin. The silk protein molecules (fibroin and spidroin in silkworms and spiders, respectively) are long polymer-like protein chains characterized by repetitive regions of typical amino acid motifs^6,10,27,28^. During the fibrillar assembly, proteins adopt a hydrogen-bonded β-sheet-rich conformation^12,26,29,30^. The reported sizes of the nano-fibrils vary between 3–300 nm in diameter^9,11,31–34^, and they appear in various fibrillated shapes, including beads-on-a-string^35–37^, uniform^31,32^, branched^36,38^, and herringbone-like^39^.

In *B. mori*, the silk gland, a tube-like organ bound by a layer of epithelial cells, is comprised of four main sections: posterior, middle, anterior, and spinneret (**Supplementary Figure S1**). The posterior section, which is shaped like a long, curled tube, is where the fibroin proteins, the main components of silk materials, are synthesized and secreted from the epithelial cells into the gland’s lumen (the inner part of the tube)^40–43^. The produced highly concentrated (∼20-30%) protein feedstock flows from the posterior to the middle section, the largest and widest part of the gland. Here, the proteins are stored in their liquid state, which preserves their native, mostly random coil fold^15,44–46^. The middle section can be further divided into posterior-middle, middle-middle, and anterior-middle parts. The epithelial cells in the middle-middle and anterior-middle sections secrete the sericin proteins ^40,47^, which later serves as the fibers’ coating layer, gluing them into a cocoon. Of note, liquid silk feedstock is highly unstable and outside the silk gland can aggregate spontaneously within seconds^48^, which represents a major limitation in the investigation of silk fiber formation mechanisms and in artificial silk fibers generation^20,49,50^. Thus, how silk proteins are stabilized inside the gland is an open scientific question that remains poorly understood.

It is in the anterior section’s long, narrow tube that the fiber-spinning process finally commences. This process is characterized by a liquid-to-solid transition (LST), during which fibroin transforms from a soluble random-coil conformation into β-sheet-rich nano-fibrils. The process ends in the anterior’s spinneret, where the silkworm’s two silk glands are connected into one tube and enters the silk press, a set of muscles that control the liquid’s compression and move it forward to the exit spigot^14,51,52^. The final microscale fiber exits as a compressed bundle of aligned nano-fibrils.

In attempts to unravel the silk fiber spinning process, the protein’s secondary structure and fibrillation mechanisms have been exhaustively studied^2,15,37,44^, mostly by examining the factors governing them. It is well established that chemical and physical forces, including shear stress, elongation flow, acidification, and metal ions, are essential for the protein’s structural transitions from storage to a fibrillated state^15,53^. For instance, while basic pH (>7) and the presence of Ca ions are associated with the stabilization of the fibroin random-coil conformation (“storage state”), more acidic pH (<6) and the presence of K, Na, Cu, and Mg ions can induce the adoption of a β-sheet conformation (fibrillation process)^54–59^. Still, the material evolution of silk in the gland consists of a more complex and dynamic chain of events that is not yet fully understood.

The most common theories describing silk feedstock behavior in the gland are the “Micellar”^60^ and “Liquid Crystal”^61,62^. Briefly, the “Micellar” model, initially described by Ji and Kaplan^60^ and further supported by several *in-vitro* studies^29,63–65^, suggests that the amphiphilic nature of fibroin facilitates the protein’s assembly into micelle-like spherical structures, which can coalesce into larger globular structures. During the spinning, shear forces align and elongate the globular-micellar clusters, initiating the liquid-to-solid transition and promoting further fibrillation. According to the “Liquid Crystal” model, introduced by Vollrath and Knight^61,62^, the silk protein feedstock inside the silk gland appears as a liquid crystalline, which allows it to flow despite its crystalline structure. In this manner, the feedstock can flow as a prealigned liquid crystal and be further aligned during the spinning. Despite the differences between the two models, they are not mutually exclusive but rather describe the different stages of silk fiber formation.

Based on multiple recent *in-vitro* studies^66–70^ performed on spidroin and spidroin-like proteins, it has become apparent that liquid-liquid phase separation (LLPS) is an essential assembly step in the silk fibrillation process. The LLPS is regulated by pH, ionic strength, and shear flow inside the silk gland. However, extracting silk feedstock from the silk gland for *in-vitro* examination often perturbs the protein’s fold and assembly state, preventing a comprehensive investigation of this phenomenon.

Indeed, the study of silk’s natural formation mechanism, in general, is challenged by the need to analyze *in-situ* the highly unstable feedstock. Most standard sample-preparation procedures involve prolonged incubation, mechanical manipulations, chemical fixation, freeze-drying, embedding, and more, which disrupt the feedstock’s native characteristics and often cause aggregation or unnatural fibrillation (**Supplementary Figure S2**).

In this study, we utilized state-of-the-art cryogenic and non-cryogenic microscopy techniques to image and characterize *in-situ* the mechanism of silk protein assembly along the whole *B. mori* silk gland (**Figure 1**), while fully preserving the natural biological environment. Our analysis revealed that fibroin, although known to have a dominantly random-coil conformation in the posterior section, assembles into micron-sized liquid spherical structures immediately after its secretion from the epithelial cells. These protein assemblies, which we term “compartments,” maintain fibroin in its soluble state as it flows through the middle section of the silk gland, thereby eliminating premature fibrillation. Fibroin undergoes several structural transformations along the silk gland in response to environmental changes. When fibroin enters the anterior section, the compartments disassemble into an unordered silk feedstock and initiate the spinning process. Imaging the entire length of the anterior section at a nano-scale unraveled a series of structural transition events in the silk feedstock. These entail the liquid alignment, formation of different types of fibrillated nano-structures, bundling of nano-fibrils, and cross-linking. Imaging the silk spinneret and spun fibers uncovered that the nano-scale organization of silk fibers is a network of cross-linked nano-bundles, each one comprised of ∼8 nm nano-fibrils, a revelation that might revise our understanding of the origins of silk fiber’s mechanics.

**Figure 1:**
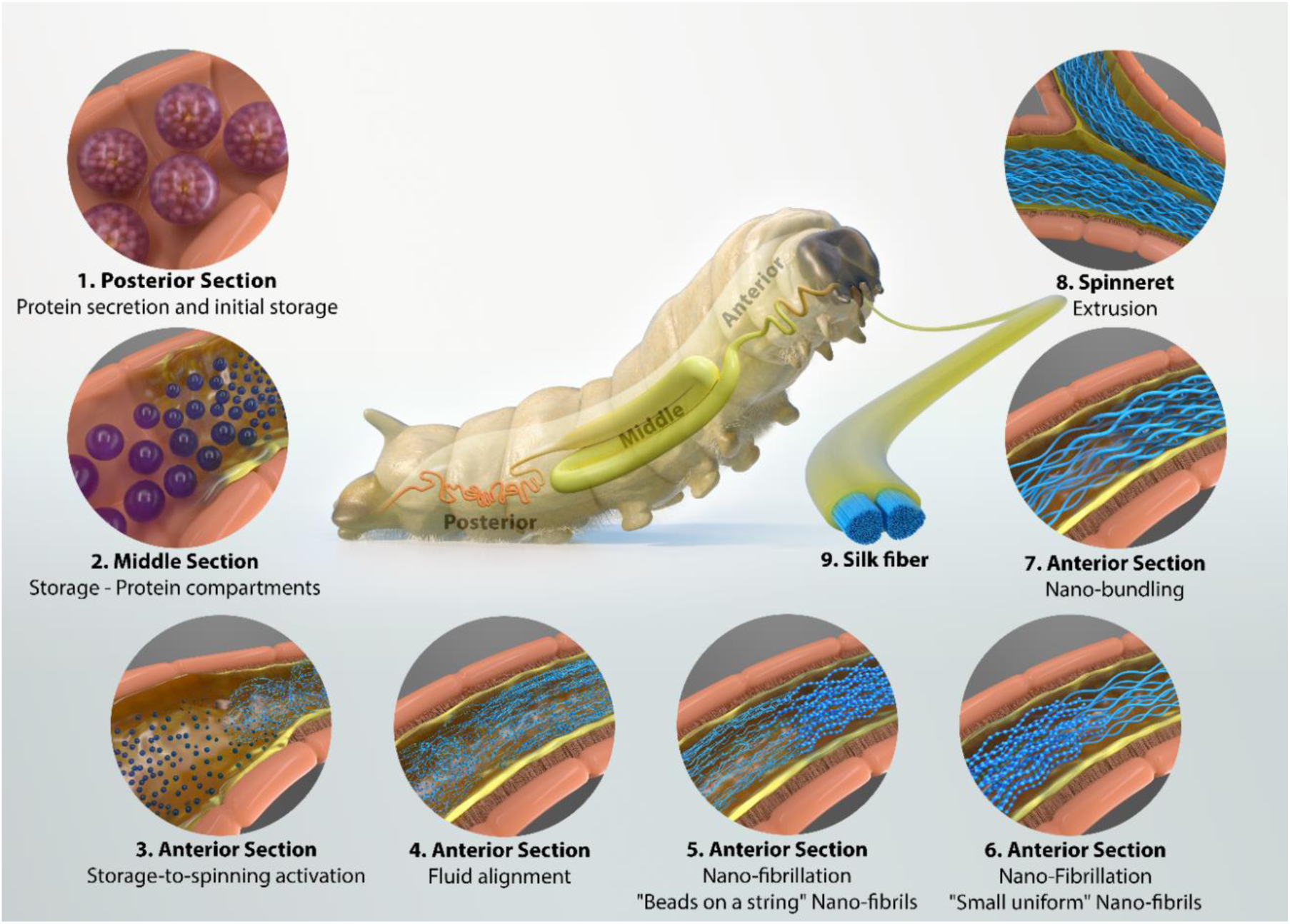
The silk spinning process. An illustration of the silk spinning process along the silk gland of a Bombyx mori silkworm: **(1)** Silk proteins (fibroin) secretion from the epithelial cells in the posterior section of the silk gland, followed by the protein initial assembly and storage inside the gland lumen. **(2)** Silk protein storage in a state of protein “compartments” in the middle sections of the silk gland. **(3)** Silk feedstock storage-to-spinning activation, accompanied by the protein compartments disassembly into an unordered fluid at the entrance to the anterior section of the silk gland. **(4-7)** The gradual structural transition and ordering of the silk feedstock in the anterior section, which includes **(4)** fluid alignment, **(5)** protein nano-fibrillation into “beads on a string” nano-fibrils followed by **(6)** their maturation into small uniform nano-fibrils which then **(7)** attach into nano-bundles. In the last part of the silk gland, called **(8)** spinneret, the two silk glands come together into a single tube as the silk feedstock is extruded and spun into the final structured **(9)** silk microfiber. The sizes of the nano-structures in the illustration are exaggerated for better visualization.

## Results and Discussion

### Storage of Silk Feedstock

To investigate the storage conditions of the silk protein feedstock inside the silk gland, we analyzed the morphology of the silk gland using confocal microscopy (**Supplementary Figure S1 f,g**). We differentiated between the gland components by staining the silk gland, while preserving its integrity, with Nile red, an environmentally sensitive dye (see **materials and methods** section). The analysis revealed that the secreted fibroin spontaneously self-assembled into micro-sized spherical structures of various sizes and inner densities (**Figure 2**). This is even though it mostly has a random-coil conformation, as revealed by Fourier transform infrared (FTIR) analysis (**Supplementary Figure S3**). Due to these spherical assemblies’ compositional heterogeneity, we will only use the general term “compartments”.

**Figure 2:**
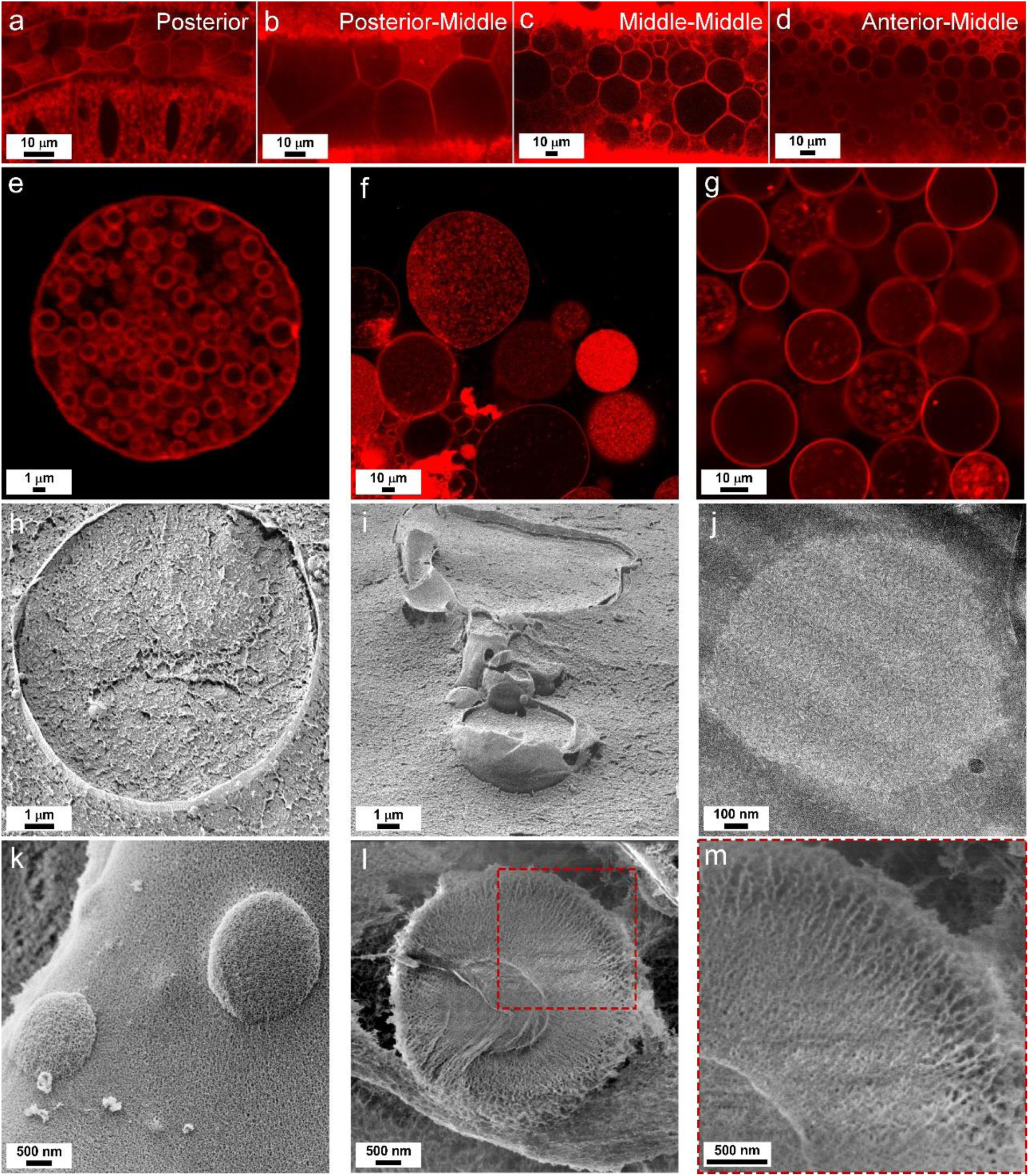
Silk protein compartments. In-situ confocal imaging of a Nile-red-stained silk gland’s (a) posterior, (b) posterior-middle, (c) middle-middle, and (d) anterior-middle parts. (e) Ex-vivo fibroin protein compartment from the posterior silk gland part and its inner structure. (f-g) Ex-vivo protein compartments from the middle parts of the silk gland. (h-i) In-situ cryo-SEM of compartments from the middle-middle part of the gland prepared by high-pressure freezing and freeze-fracture. (j) A TEM image of a thin section obtained from the middle-middle part of the gland prepared by high-pressure freezing and freeze-substitution. (k-m) In-situ cryo-SEM images of compartments from the beginning of the anterior part of the silk gland prepared by high-pressure freezing and freeze-fracture. (m) enlarged section of the image (l) highlighted with a red square. Scale bars are shown in the left bottom corner of each image.

We found that in the posterior region of the silk gland (**Supplementary Figure S1 e,f**), the lumen (inner part of the gland tube) is densely laden with protein compartments (**Figure 2 a** and **Supplementary Figure S4 a-d**). These compartments are tightly packed with small spherical protein assemblies that fill their entire interior (**Figure 2 e** and **Supplementary Figure S4 d-i**). The diameter of the inner spherical assemblies ranges from a few hundred nanometers to two micrometers (**Figure 2 e, Supplementary Figure S4 e-h** and **Supplementary movies 1** and **2**). Real-time confocal imaging revealed that these small spherical assemblies move within the larger micro-sized compartment (similar to protein colloidosomes^71^), (**Supplementary movie 3**).

Our assumption is that the process of fibroin secretion from the gland’s epithelial cells to its lumen is responsible for the formation of the small spherical assemblies. In this process, Golgi vesicles transport the synthesized fibroin protein inside the epithelial cells at a highly concentrated state and secrete it into the lumen in the form of liquid “spherical masses” lacking the Golgi lipid component^13,43,72^. This assumption is corroborated by the observation that the small spherical assemblies encapsulated inside the compartments are present only in the posterior section and not in the middle section. While flowing into the middle section of the silk gland, the spherical assemblies disassemble, forming a uniform phase inside the compartment.

We attribute this colloidosome-like assembly to the “Micellar” theory, ^29,63–65^, which describes the assembly of fibroin protein into small (nano-scale) micelles that then merge into larger colloidosomal globules (**Figure 2 e** and **Supplementary Figure S4 e**). However, the “Micellar” theory claims an internal micelle-like organization of the fibroin protein chains, which we could not confirm *in-situ,* because the inner structure of the compartments dynamically changing as they “mature” in the middle section of the silk gland. Such changes, for example, manifested by the disappearance of the spherical assemblies inside the compartments (**Figure 2 b-d, f, g**, and **Supplementary Figure S5**). This observed disappearance eliminates the possibility that the spherical assemblies take part in forming the nano-fibrils at the latter stages of silk fibrillation.

In the middle section of the silk gland, the size of the compartments varies greatly, from 200 nm to 50 um in diameter, filling the inner volume of the gland (**Figure 2 b-d, f, g**). The largest compartments (15–50 um in diameter) are found in the posterior-middle part, densely packing the glands’ lumen (**Figure 2 b** and **Supplementary Figure S5 a-c**). The compartments gradually shrink as they flow into the middle-middle (**Figure 2 c** and **Supplementary Figure S5 d-e**) and anterior-middle (**Supplementary Figure 2 d** and **Supplementary Figure S7**) sections. Concurrently, there is a gradual decrease in the feedstock’s pH from 7–7.5 to 5.5–6.5^55,63,73,74^, and changes in the metal ion composition^58,75,76^, along the silk gland, which likely causes the compartments’ destabilization. The compartments have an outer shell, visible by Nile-red staining, but no distinct inner structure (**Figure 2 g** and **Supplementary Figure S5 f-i**); they contain an unstained material or freely moving aggregates (100–300 nm) (**Figure 2 f-g** and **Supplementary movie 3**). The difference in Nile-red staining, which exhibits polarity-sensitive fluorescence^117, 118^, possibly indicates that the proteins at the outer shell of the compartments have a more arranged and densely packed structure compared to less structured proteins at the compartment core.

Cryogenic scanning electron microscopy (Cryo-SEM) images of freeze-fractured compartments from the middle-middle gland (**Figure 2 h,i** and **Supplementary Figure S6 a-d**) provided further support for the results of the confocal microscopy analysis. The micro-sized spherical assembly consists of a well-defined shell, coating an unstructured inner part. This distinct difference suggests that the fibroin proteins assemble differently in these two regions and probably adopt separate conformations.

Interestingly, a transmission electron microscopy (TEM) analysis of the middle-middle gland region via freeze-substitution followed by ultra-microtome sectioning (see **materials and methods** section) revealed that silk compartments assume a dense inner phase, similarly to biomolecular condensates^77,78^, surrounded by a less concentrated continuous phase (**Figure 2 j** and **Supplementary Figure S6 e,f**). This suggests that the internal organization of the compartment is dynamic and tends to assemble and disassemble while inside the silk gland.

To probe the compartments’ stability outside the silk gland, we placed extracted compartments into a diluted environment (milli-Q water or PBS). We observed that compartments are stable for a few hours up to two days, and then shrink and disappear or fibrillate, indicating that the movement of individual protein molecules between concentrated and diluted phases is relatively slow. Thus, the compartment’s characteristics enable protein stabilization and storage inside the silk gland.

As the feedstock enters the gland’s anterior section, the compartments’ morphology changes, as gleaned from our cryo-SEM analysis (**Figure 2 k** and **Supplementary Figure S8**), and their size drastically decreases to an average of 1.3 μm in diameter (**Supplementary Figure S8 h**). The shell of these compartments also changes, becoming less defined and resembling more the unstructured bulk protein surrounding it (**Figure 2 k** and **Supplementary Figure S8 a-d**). Freeze-fractured compartments lack a clear outer boundary but possess a dense core surrounded by elongated and radially-oriented chains (**Figure 2 l,m** and **Supplementary Figure S8 e-g**). This radially-oriented structure resembles an oversized micelle, but given its size (1–2.5 um), it is not composed of a single layer of molecules. Thus, the proteins must assemble into chains that interact to form such a spherical microstructure. This nano-scale organisation might result from the process of compartments dissasembly and the protein’s transition from its storage state to an active processing state.

### Feedstock Storage-to-Spinning Transition

Our further cryo-SEM analysis of the freeze-fractured anterior sections of the silk gland showed that once the silk feedstock enters the anterior region, not only the size of the protein compartments reduces, but also their quantity (**Supplementary Figure S8 a-d**). We further observed that the liquid silk proteins are phase-separated into a compartment state and a surrounding bulk protein-containing fluid. Both states co-exist at the beginning of the anterior section of the gland, but as the feedstock continues to flow towards the spinneret, the compartments dissociate into an unstructured fluid.

This dissociation is brought about, at least in part, by changes in pH and ion levels. According to literature reports, the fluid’s flow from the middle to the anterior section is accompanied by a drop in pH from ∼7, a level that endows the silk feedstock with more stability and less sensitivity to shear, to ∼ 5–6^15,55,73^, which makes the feedstock more shear-sensitive, and the fibroin protein more prone to fibrillation^55,74,79^. We recently showed that this transition is also accompanied by a drastic depletion of metal ions from the fibroin feedstock^80^. The ions diffuse out of the silk feedstock to the sericin phase and epithelial cells^80^. By triggering the compartments’ disassembly, the alterations in pH levels and metal ions lead to the domination of the “unstructured fluid” state. This “unstructured fluid state” occupies a substantial length of the first part of the anterior section.

### Fluid Alignment

The increase in flow velocity and shear rate due to the sharp decrease in the gland’s diameter at the beginning of the anterior section^81^ is expected to cause the unstructured fluid to align with the flow’s direction. This alignment is clearly seen in our cryogenic electron microscopy analysis (**Figure 3 a** and **Supplementary Figure S9**). Fibroin solution is a non-Newtonian shear-thinning fluid, and its molecules, which are long polymer-like chains, unfold and align upon applied shear flow (similarly to the behavior of other polymers^82–84^), reducing the overall fluid viscosity^74,85–87^. The flow at this point only induces the proteins’ reorganization and alignment (not fibrillation), starting either from the edges of the inner gland’s tube or from the center of the fluid and spreading to the rest of the material (**Supplementary Figure S9**). Thus, at this stage, the conditions applied to the feedstock do not reach the critical shear rate, shear stress, and accumulated energy sufficient to initiate fibrillation^76,88,89^. The proper stretching and aligning of the protein chains is a crucial stage for their fibrillation, as this ensures the nano-fibrils form in a highly oriented manner ^76,80,85,90^. Interestingly, TEM imaging of a freeze-substituted and microtome-sectioned anterior gland (**Supplementary Figure S10**) revealed a non-homogeneous phase separation in the form of loosely packed, unconnected nano-scale assemblies surrounded by densely packed, aligned fluid (**Supplementary Figure S10 a,b**). The size of these hexagon-shaped nano-scale assemblies, 20–30 nm (**Supplementary Figure S10 c-f**), is comparable to that of fibroin monomers, as measured in *in-vitro* studies^35,37^.

**Figure 3:**
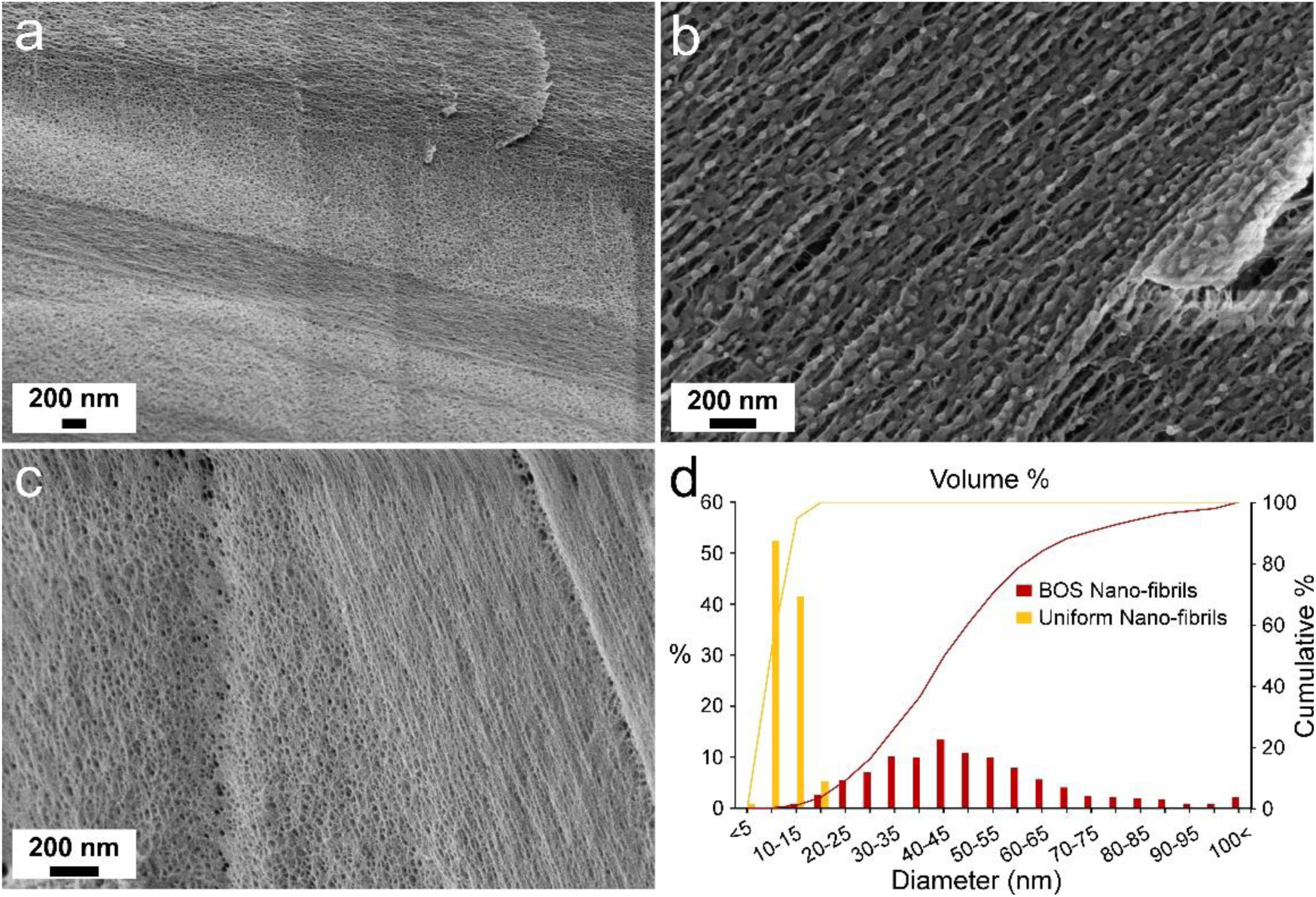
Silk Nano-fibrils. In-situ cryo-SEM of silk feedstock in different parts at the middle region of the anterior section of the silk gland prepared by high-pressure freezing and freeze-fracture showing **(a)** the protein liquid alignment, **(b)** the formation of “beads-on-a-string” nano-fibrils, and **(c)** the small-uniform nano-fibrils. **(d)** Size distribution of the nano-fibrils in terms of volume percent. Size measurements were produced by image analysis. Scale bars for **(a-c)** are shown at the left bottom corner of each image.

### Fibrillation

As the feedstock flows inside the anterior section, the stretched and aligned silk protein chains interact and start to fibrillate, forming elongated, aligned nano-fibrils (**Figure 3 b,c**). Our microscopy analysis revealed the presence of two distinct fibrillated species inside the silk gland anterior: nano-fibrils with a “beads-on-a-string” (BOS) shape (**Figure 3 b** and **Supplementary Figure S11**) and ones with a uniform shape (**Figure 3 c** and **Supplementary Figure S12**).

The nano-fibrils’ size distribution and average diameter were determined via automated image analysis of the cryo-SEM images (see **materials and methods** section). The BOS nano-fibrils were found to be a long, fibrillated species (**Figure 3 b** and **Supplementary Figure S11**) that are decorated with spherical “beads” along its entire length. Their average diameter is 33.94±15.36 nm (assuming a cylindrical shape), but they broadly vary in size (**Figure 3 d**), with 96% of the nano-fibril volume composed of nano-fibrils with a diameter greater than 20 nm.

Similar BOS nano-fibrils were reported to form *in-vitro* by reconstituted silk fibroin (RSF)^2,35,36,91^. The RSF BOS nano-fibrils had a characteristic diameter of 20-30 nm and were suggested to be composed of aligned, connected monomers or a cluster of hydrophobic domains of several fibroin molecules. Recently, we showed, using nano-Fourier transform infrared spectroscopy, that the “beads” decorating the nano-fibrils in the silk gland act as an intermediate state, or “nanocompartments”, and that they take part in the protein conformational transition from a native protein fold into an aggregated β-sheet conformation^37^.

The uniformly shaped nano-fibrils are aligned and are smaller in size than the BOS nano-fibrils (**Figure 3 c** and **Supplementary Figure S12**), featuring a diameter of 8.27±2.65 nm. A size analysis revealed that most (∼94%) of the small-uniform nano-fibrils’ volume is composed of fibrils with a diameter of 5–15 nm (**Figure 3 d**). These nano-fibrils are the smallest structured unit identified in the material, serving as the basic building blocks of the silk fiber. Such small diameters of nano-fibrils have not been previously reported for *B. mori* fibroin^2,11,35,92^.

Most studies of natural fibroin nano-fibrils assume that the smallest nano-fibrillar structural unit in *B. mori* silk fibers has a diameter range of 20–100 nm^11,31,33,91,93^. The discrepancy between these reports and our results may be due to the way by which the nano-fibrils were generated *in-vitro,* their ability to form transient nano-fibrillar BOS shapes, or the limited capability to differentiate between nano-structures in solid silk fibers.

Both types of nano-fibrillar species were observed to form separately and independently from the unstructured feedstock. The BOS nano-fibrils are partially composed of smaller nano-fibrils and, upon strain (caused by sample manipulation and sublimation), can be stretched to form small-uniform nano-fibrils (**Supplementary Figure S13**). We conclude that BOS nano-fibrils are a less mature fibrillated species that transitions into small-uniform nano-fibrils as a result of the stretching caused by the elongational shear flow. However, they are not essential for the formation of small-uniform nano-fibrils.

The silk feedstock continues to flow even when the proteins are already in their fibrillated state (structured fluid). The silk feedstock fibrillation increases the viscosity and overall resistance to fluid deformation^85,94^. The flow of the feedstock possibly causes nano-fibrils to slide on top of one another, the result of which is extremely high shear.

### Nano-Bundle Formation

We further observed that the nano-fibrils formed in the anterior section intertwine to form “nano-bundles” (**Figure 4**), which are cross-linked by small connecting fibril bridges. The structures are initially gradually distributed from the inner wall of the silk gland tube, where the shear stress is greatest^81^, toward the center of the tube, which was less populated with “nano-bundles” (**Figure 4 a,b**, and **Supplementary Figure S14**). Eventually, the entire feedstock inside the lumen is structured as cross-linked and nano-bundled morphologies. An image analysis of the newly formed nano-bundles (**Figure 4 c**) found the nano-bundles to be 15.87±4.64 nm in diameter. The volume of the analyzed fibrillated species (**Figure 4 e**) comprised ∼95% of fibrils with a diameter larger than 10 nm, indicating that most of the material formed nano-bundles, with only ∼5% of the analyzed fibrils remaining as unbundled nano-fibrils (<10 nm in diameter).

**Figure 4:**
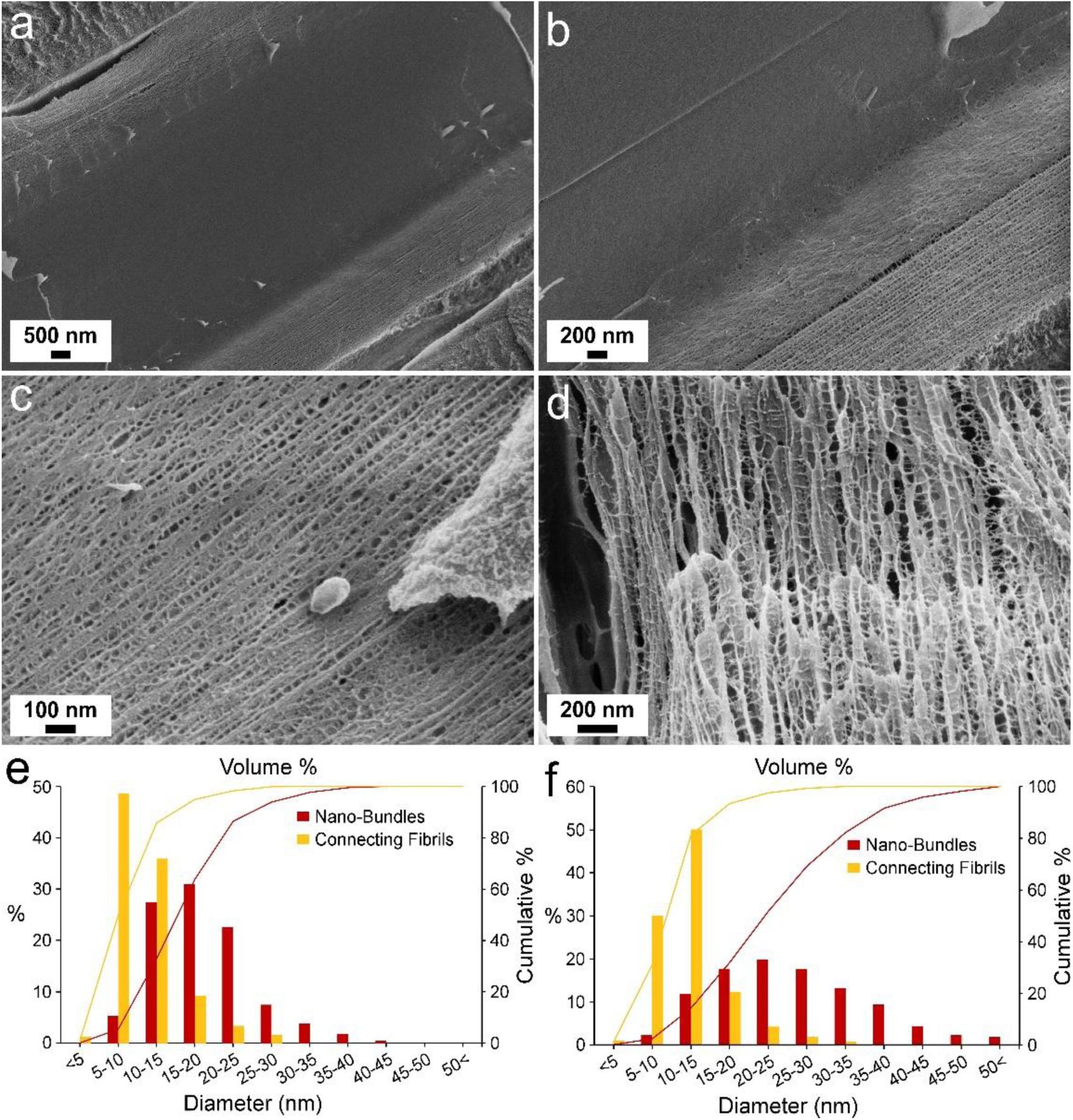
Silk Nano-Bundles and cross-linked structure. In-situ cryo-SEM images of silk feedstock in the last parts of the anterior section of the silk gland, prepared by high-pressure freezing and freeze-fracture. **(a,b)** Nano-bundles gradually form inside the gland. **(c)** Newly-formed nano-bundles. **(d)** Further-sheared nano-bundles. Size distribution of **(e)** newly-formed and **(f)** further-sheared nano-bundles and the connecting nano-fibrils showing the volume percent of fibrils with different diameters. Size measurements were produced by image analysis. Scale bars for **(a-d)** are shown at the bottom left corner of each image.

A size analysis of the fibril bridges connecting the nano-bundles (**Figure 4 c** and **Supplementary Figure S14 h,i**) found them to be 8.33±3.24 nm in diameter, similar to that of the small-uniform nano-fibrils (∼8 nm), the basic building blocks of the material (**Figure 3 c**). The volume of the analyzed fibril bridges mostly consisted (∼86%) of fibrils with diameters smaller than 15 nm (**Figure 4 e**). We concluded that the nano-bundles are linked by the same basic nano-fibrils that construct them. Our assumption is that a single nano-fibril can start in one nano-bundle and end in another, thus linking the two together.

As the nano-bundles propagate along the gland tube towards the spinneret, the width of the nano-bundles tends to increase. We were able to visualize the event of “shearing” by imaging a defective part of the gland that caused the feedstock to increase its flow, thus increasing the shear during the preparation of the freeze-fractured anterior section (**Figure 4 d** and **Supplementary Figure S15**). In this section of the anterior, the nano-bundles’ average diameter increased to 19.42±7.76 nm. The developed fibrillated feedstock volume comprised ∼98% nano-bundles with a diameter larger than 10 nm, and only ∼2% were unbundled nano-fibrils with less than 10 nm diameter (**Figure 4 f**). At this point, each nano-bundle’s volume (assuming it has a cylindrical shape) equals the volume of more than five small-uniform nano-fibrils.

This network structure of cross-linked nano-bundles is especially interesting. It shed light on the hierarchical structure of the final silk fibers. Until now, the most common model for describing the silk fiber’s structure has been a micro-sized bundle consisting of aligned nano-fibrils woven together by weak physical interactions^2,31^. Here, we show that the material has an additional hierarchical stage of nano-bundles linked by connecting fibril bridges. The cross-linking of the material reduces pull-out of nano-fibrils upon applied strain and inhibits fracture propagation, a phenomenon that explains silk fibers’ high strength and toughness.

### Silk Microfiber Formation

We used Nile-red staining and confocal imaging to visualize the spinneret, the last part of the silk gland, where the nano-bundled, cross-linked silk feedstock is spun into the final fiber (**Figure 5 a** and **Supplementary fFigure S16 a-e**). At the spinneret, the silkworm’s two silk glands merge to form a single tube just before entering the silk press (see confocal images of the spinneret in **Supplementary Figure S16 d,e** and light microscopy images of the spinneret ultra-microtome sections prepared for TEM imaging in **Supplementary Figure S16 f,g)**. The two separate feedstocks appear together inside the spinneret, forming a bave of two silk fibers glued by sericin as they exit from the spigot.

**Figure 5:**
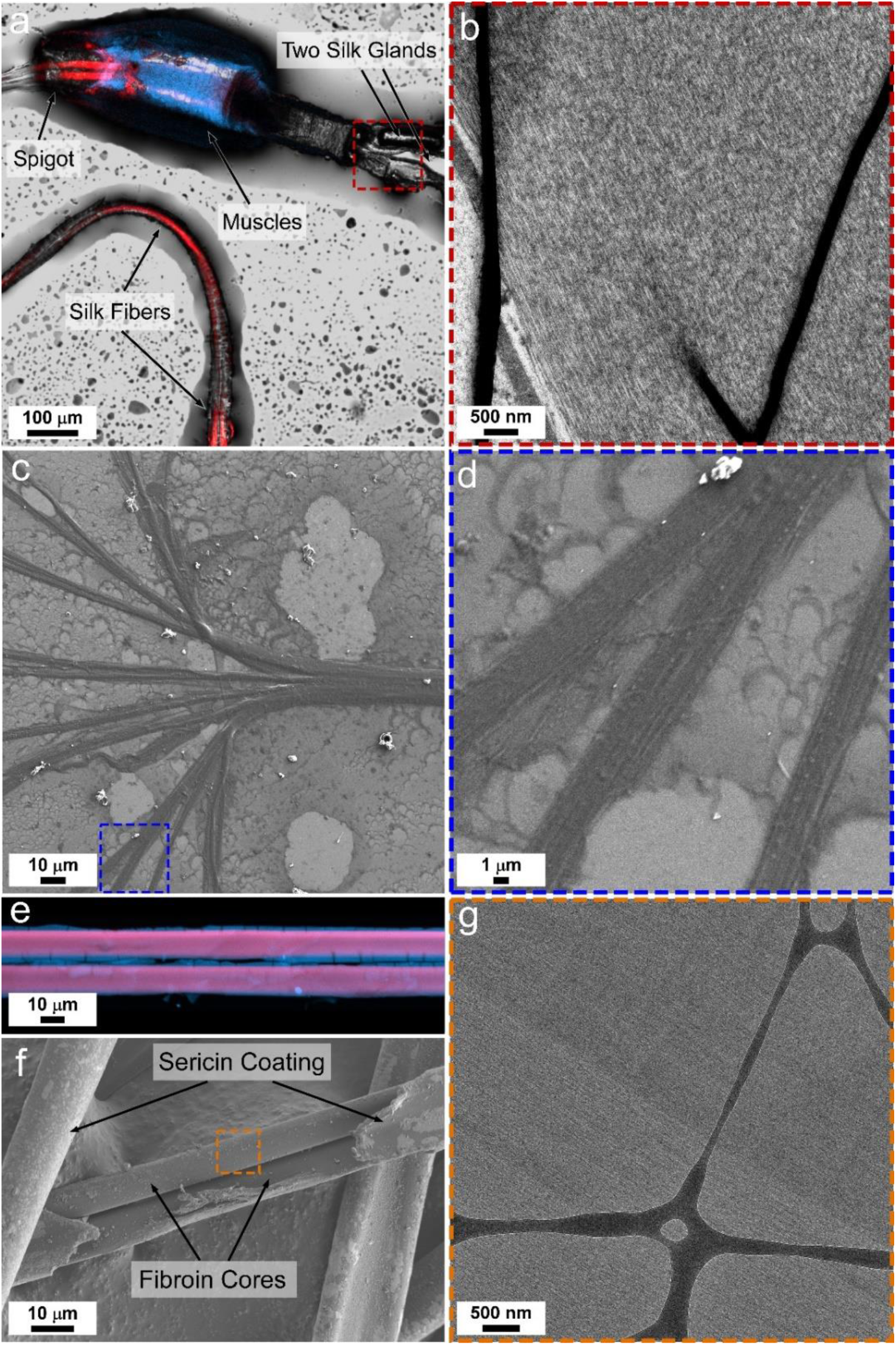
Silk fiber Formation. **(a)** Confocal image of a silk gland spinneret, including the two silk glands’ intersection, the silk gland’s press muscles (blue), exit spigot, and spun fibers (red). **(b)** A TEM image of a thin section from the intersection of the two silk glands (corresponding to the area marked by a red square in **(a)**) showing the feedstock fibrillated structure just before the final fiber formation, prepared by high-pressure freezing and freeze-substitution. **(c)** SEM image of silk feedstock starting to form a silk fiber. **(d)** Magnification of the area marked by a blue square in c. **(e)** Confocal microscopy image of fully-formed silk fibers. **(f)** SEM image of fully-formed silk fibers. **(g)** A TEM image of a thin section of silk fiber embedded in EPON (corresponding to the area marked by an orange square in **(f)**). Scale bars are shown at the bottom left corner of each image.

The entrance to the spinneret was successfully imaged by TEM (**Figure 5 b** and **supplementary Figure S17**) after freeze-substitution and ultra-microtome sectioning (light images in **Supplementary Figure S16 f,g)**. While flowing in the spinneret, the feedstock maintains a dense and aligned fibrillated composition. Since the dense and crowded structure of the feedstock limit the TEM image analysis, manual measurements of nano-fibrillated structures’ sizes were performed. Similarly to the cryo-SEM images, the average diameter of the unbundled nano-fibrils and nano-bundles was 8.07±1.14 nm and 16.05±4.54 nm, respectively (the average diameter of the total measured nano-fibrillated structures was 13.44±5.34 nm). These results support our previous conclusions from the cryo-SEM images, that the material is constructed from small unbundled nano-fibrils, the basic building blocks, and larger nano-bundles.

The structured liquid feedstock dehydrates and forms a solid silk fiber upon exiting the gland’s spigot. By allowing the silk feedstock to exit from the silk gland into water and stick on a glass surface, we were able to capture the final step of the spinning. The obtained structure was a not-fully-bundled fibroin fiber (**Supplementary Figure S18**) that lacked the sericin coating, thus exposing its structure (**Figure 5 c,d**). SEM imaging revealed the silk fiber to be a bundle of highly aligned, stretched nano-fibrils (**Supplementary Figure S19**). The arrangement and nano-structure of the fiber are better seen when SEM imaging penetrates to its depth (**Supplementary Figure S19 a,c,e**) compared to its outer surface (**Supplementary Figure S19 b,d,f**). In addition, some spherical structures are observed on the surface of the fiber (**Supplementary Figure S19 c,d**). These might be fibroin protein compartments spilled out from more posterior sections of the silk gland during the sample preparation process or not-yet-fibrillated proteins left in the feedstock.

Using atomic force microscopy (AFM), we were able to image the fiber’s surface topography (**Supplementary Figure S20**), including the bundle structure and the nano-fibrils forming it. The high roughness of the fiber (**Supplementary Figure S20 f,g**) is a result of the unfinished final spinning stage of dehydration and stretching. Thus, some nano-fibrils or small chunks of bundles were not firmly attached to the fiber and separated from it. The disentangled fibrils or small bundles are attached to the surface next to the larger fibers (**Supplementary Figures S19 c,d**, and **S20 d-g**).

The size of the imaged fibrils constructing the fiber and the ones coming out of it roughly varies from a few tens of nm to hundreds of nm in diameter. The measured size of the nano-fibrillated structures aligns with previous reports stating that silk nano-fibrils are 20–300 nm in diameter^9,11,31–34^. Although the fiber is partially disentangled, it is hard to distinguish between nano-fibrils and measure their exact size due to the density of the material. Furthermore, we assume that most of the fibrils imaged on the surface of silk fibers or those detached from them are nano-bundles. Some of them are the nano-bundles that, as we observed, constructed the silk feedstock at the final stage, just before exiting the silk gland (**Figure 4**), and some are larger chunks of bundles made of fibrillated material.

The final native silk fibers form as a bave of two fibroin fibers coated by a sericin layer (**Figure 5 e,f**). TEM images of native fibroin fibers after the fibers’ ultra-microtome sectioning (**Figure 5 g** and **Supplementary Figure S21**) revealed their highly aligned, tightly packed nano-fibrillated structures. The measured diameter of the nano-fibrils was found to be in the range of 10–25 nm. It should be emphasized that silk fibers’ highly dense structure permits accurate measurements of the fiber’s nano-structure by using image analysis. This diameter range is in agreement with that of previously observed nano-bundles in silk feedstock.

## Conclusions

In conclusion, silk’s natural material evolution, from its stable storage as liquid feedstock to its transition into a highly ordered solid fiber, is a carefully tuned, multi-step process involving several events of phase separation and macromolecular assembly. Here, we unraveled the actual process by *in-situ* imaging along the entire silk gland while preserving the silk feedstock’s natural unaltered state. We show how the protein assembles at each stage to dictate the material properties and serve the needs of each processing phase.

Although the protein was believed to be mostly unstructured during storage inside the posterior and middle sections of the silk gland, it assembles into micron-sized compartments, which are believed to suppress premature fibrillation and unwanted aggregation of the highly shear-sensitive feedstock. The protein compartments undergo several structural transitions along the gland, which are yet to be fully understood, but are nevertheless indicative of the compartments’ flexibility and adaptable properties for different purposes. These include a platform for correctly folding newly secreted proteins, stabilizing proteins during storage, and controlled disassembly before fibrillation. Notably, the compartments do not take part in the fibrillation inside the gland, as they disassemble just before the feedstock starts the structural transition. Under the action of shear and flow, the unordered protein feedstock aligns and fibrillates in the narrow anterior section. In this process, the protein assembles into two fibrillated species: less mature “beads-on-a-string” nano-fibrils (∼30 nm) and smaller uniformly-shaped nano-fibrils (∼8 nm). While both species appear to form independently, the stretch or shear of the former nano-fibrils results in their transition into the small-uniform ones, which serve as the fundamental building blocks of the material. Importantly, the last structure maturation of the feedstock, before it is spun into a solid fiber, is the development of nano-bundles cross-linked by “bridges” of nano-fibrils. Hence, the final silk fiber comprises densely packed and cross-linked nano-bundles (∼20 nm), a revelation that will help better understand its mechanical properties.

Thus, our results shed light on long-lasting mysteries and theories regarding natural silk fiber formation, broaden our understanding, and provide actual validations and images of this unique process. We believe that the reported finding is relevant not only for the generation of *B.mori* silk fiber, but also represents a general route of silk feedstock processing used by various organisms, including spiders. The findings will help mimic the process, design better biomaterials, and develop new concepts for improved material processing and synthesis.

## Materials and Methods

### Silk Gland Extraction

All *Bombyx mori* silkworms used in the study were grown in the lab at a constant temperature of 25–26°C and fed freshly picked mulberry leaves. Before dissection, silkworms were anesthetized with N2 for 20 min (for humane reasons and to prevent damage to the silk gland during dissection). Silkworms were dissected using fine scissors and razor blades by applying a longitudinal dorsal incision (without removing the head) and fixing the skin to a surface with needles. The stomach of the silkworm was removed, and the tissue was gently washed with Milli-Q water. The two silk glands were gently detached in their entirety from the rest of the tissues, with the silkworm’s head still attached, and gently rinsed with PBS buffer.

### Silk Gland Imaging

#### Confocal Imaging of Silk Glands

Prior to imaging in the confocal microscope, the extracted silk glands were incubated in a phosphate buffered saline (PBS) solution with Nile red at a final concentration of 3 μM, for 2–3 hours, in a 35 mm μ-dish with a glass coverslip bottom (Ibidi, USA). Imaging of the anterior section and spinneret was done on silk glands extracted from fifth instar silkworms just after starting to produce silk for constructing the cocoon. Imaging the entire silk gland, specifically the middle and posterior sections, was performed on silk glands extracted from 8–12-day-old silkworms after a very gentle dissection and cleaning procedure. Young silkworms were used to overcome the limited ability to image the larger parts of the silk gland (middle and posterior sections) due to their thick layer of epithelial cells.

Two confocal systems were used for imaging:

- Zeiss LSM 800 confocal system connected to an Axio Observer inverted microscope (Carl Zeiss AG, Germany). Imaging was performed with a PL APO 20x/0.8 M27 (FWD=0.55mm) objective lens. An excitation laser of 561 nm and an emission detection range of 595–700 nm were used for Nile-red fluorescence acquisition. An excitation laser of 405 nm and an emission detection range of 410–546 nm were used for intrinsic fluorescence and green Nile-red fluorescence acquisition.
- Dragonfly spinning disc confocal system (Andor Technology Ltd., Belfast, UK), connected to a Leica Dmi8 microscope (Inverted) (Leica GMBH), CF40 disk. Slides overview was performed with HC PL APO 20x/0.75 CS2 air objective lens (stitched tiles images with 10% overlap). Imaging was performed with HC PL APO 40x/1.10 W CORR water objective lens. An excitation laser of 561 nm and 600/50 nm emission filter were used for Nile red fluorescence acquisition.

The 3D images were reconstructed using Imaris software.

### High-pressure Freezing of Silk Glands

For electron microscopy imaging of the anterior section of the silk gland, fifth instar silkworms were dissected just after starting to produce silk for constructing the cocoon. The silk gland’s anterior section was briefly cut to posterior and anterior regions and placed on aluminum discs (Wohlwend Engineering Office, Switzerland), as seen in **Supplementary Figure S22 a-c**. Samples of the middle section of the silk gland were prepared by extracting silk glands from 8–12-day-old silkworms and gently placing them in aluminum discs. The samples were then covered with a second disc, immediately frozen in an EM ICE high-pressure freezing (HPF) machine (Leica Microsystems, Vienna, Austria), and kept in liquid nitrogen for future processing.

The samples intended for cryo-SEM imaging (after a freeze-fracture procedure) were sandwiched between two aluminum discs with a depression depth of 100 µm. The surface of the discs was scratched with a razor blade to prevent the samples from separating from the discs during the freeze-fracture process. Empty spaces between the sample and the disc walls were filled with PBS buffer to avoid air bubbles. The samples intended for TEM imaging (after a freeze-substitution procedure) were placed on an aluminum disc with a depression depth of 200 µm and covered with a flat disc.

### Cryo-SEM Imaging of High-Pressure Freezing and Freeze-Fractured Silk Glands

High-pressure frozen samples were mounted on a holder and transferred to a BAF 60 freeze-fracture device (Leica Microsystems, Vienna, Austria) using a VCT 100 shuttle (Vacuum Cryo Transfer Device, Leica). The samples were freeze-fractured at a temperature of -120°C under a pressure of about 5x10^-6^ mbar by rapidly removing the upper disc using the instrument’s built-in knife. Samples were transferred to an Ultra 55 SEM (Zeiss, Germany) using the VCT 100 shuttle and observed using a secondary electron InLens detector at an acceleration voltage of 1-2 kV, temperature of -120°C and pressure of about 5x10^-6^ mbar (**Supplementary Figure S22 d,e)**. Some structures could only be revealed after sublimation (etching) at (-100)–(-90)°C for 10–15 min.

### TEM Imaging of High-Pressure Freezing and Freeze-Substituted Silk Gland Sections

High-pressure frozen samples were freeze-substituted in an AFS2 freeze-substitution device (Leica Microsystems, Austria) in anhydrous acetone containing 1% glutaraldehyde and 0.2% tannic acid. After 60 hours of incubation at -90°C, samples were warmed up to -20°C over 24 hours, and washed three times with acetone. The samples were then incubated for 4 hours in acetone containing 2% osmium tetroxide and 0.2% uranyl acetate, followed by another washing step with acetone. Next, samples were brought to room temperature and then infiltrated for five days in a series of increasing concentrations of Agar 100 epoxy resin in acetone (**Supplementary Figure S22 f-h**). After epoxy polymerization at 60°C, 60–80 nm sections were cut using a UC7 ultra-microtome (Leica Microsystems, Vienna, Austria), stained with lead citrate, and examined in a Tecnai T12 transmission electron microscope (Thermo Fisher Scientific, Eindhoven, The Netherlands) operating at 120 kV. Images were acquired using a TemCam-XF416 4k x 4k CMOS camera (TVIPS).

### Image Analysis

All electron microscopy images were analyzed with open-source software, Fiji^95^, and Ilastik^96^. Size analysis of silk nano-fibrils and nano-bundles was done using semi-automated image analysis. A detailed explanation, the code used for the analysis, and the trained Ilastik models can be found on our GitHub (https://github.com/WIS-MICC-CellObservatory/Silk-threads-analysis). Below, we provide the main steps taken for the analysis.

#### Developing identification/segmentation model

We trained an auto-context Ilastik model to identify and segment the fibrillated structures in an image. To optimize the identification, four independent models were developed for each type of fibrillated structure: Beads-on-a-string (“BOS”) nano-fibrils, uniform nano-fibrils (“stretched”), newly-formed nano-bundles (“inside”) and further-sheared nano-bundles (“outside”). For each model, the training used at least three representative images.

#### Fibrillated structure identification

Using Fiji script, the identified fibrillated structures in the image were first skeletonized and then segmented by removing the skeleton intersection points. Each segment was then identified as a “main fiber” (in the direction of the flow) or “connecting fiber” (roughly perpendicular to the direction of the flow) according to its orientation. Further analysis was done to get general information such as mean local thickness^97^ and length.

#### Semi-automated analysis

Fibrillated structures were analyzed to retrieve information regarding their thickness and distance from one another. Multiple images (between 3–6) were analyzed for each type of structure. The average diameter of the fibrillated structures was calculated by averaging all the “thickness” values obtained along the fibrillated structures, assuming they have a cylindrical structure. The volumetric percent of the size distribution was obtained by calculating the theoretical volume at each measured pixel, assuming a cylindrical structure, and then dividing the cumulative volume of fibrils at a given size range by the total volume.

In addition, non-automated size measurements of other types of images were done manually using simple FIJI measurement tools.

### Native Silk Fiber and Fiber Formation Imaging

Native silk fibers were carefully removed from the silk cocoon, placed straight on a glass slide, and then glued at the edges by tape. The fibers were stained overnight in 3 μM Nile-red solution, covered, and sealed with a cover slide.

The fiber’s formation sample was prepared from the most matured silk feedstock, just before it was spun into a fiber. A silk gland extracted from an 8–12-days-old silkworm was gently placed in a 35 mm μ-dish with a glass coverslip bottom (Ibidi, USA). PBS solution with Nile red at a final concentration of 3 μM was placed on top of the silk gland, and the most anterior part of the gland was briefly cut with a razor blade. The gland was incubated for 3 hours to allow the feedstock to flow out of the gland. Most of the buffer solution was removed; care was taken not to affect the extracted silk feedstock and to allow it to stick to the glass surface.

### Confocal Imaging of Native Silk Fibers

Imaging was performed using a Zeiss LSM 800 confocal system (Carl Zeiss AG, Germany) with a PL APO 20x/0.8 M27 (FWD=0.55mm) objective lens. An excitation laser of 561 nm and an emission detection range of 595–700 nm were used for Nile-red fluorescence acquisition. An excitation laser of 405 nm and an emission detection range of 410–546 nm were used for intrinsic fluorescence acquisition.

### Atomic Force Microscopy Imaging of Fibers

Samples were imaged on an AFM (JPK Nano wizard 4 AFM, Germany), using AC240 cantilever in AC mode at 23°C and 35% humidity. Images and 3D reconstructions were produced using Gwyddion software^98^.

### SEM Imaging of Fibers

Imaging was acquired using an InLens detector at an acceleration voltage of 1–1.5 kV and an SE2 detector at an acceleration voltage of 3 kV.

### TEM Imaging of Silk Fiber Sections

Native silk fibers, straightly placed on a glass slide, were fixed in Agar 100 epoxy by polymerization at 60°C. Fiber sections 60–80 nm in length were cut using a UC7 ultra-microtome (Leica Microsystems, Vienna, Austria). Imaging was performed at liquid nitrogen temperature conditions with a Titan Krios G3i TEM (Thermo Fisher Scientific) operated at 300 kV. Specimens were introduced into the Titan Krios via an Autoloader, and images were recorded with Gatan K3 Direct Electron Detector used in electron counting mode. The electron fluence used was 20 e-/Å2.

#### Protein Secondary Structure Analysis Using Fourier Transform Infrared Spectroscopy

Protein structural analysis of degummed silk fibers and native silk feedstock extracted from the posterior middle section of the silk gland (as previously described) was performed using FTIR spectrometer Nicolet iS50 with an introduced Attenuated Total Reflection (ATR) accessory (Thermo Scientific). The atmospheric compensation spectrum was subtracted from the original FTIR spectra, and a second derivative was applied for further analysis. To resolve the spectra, OMNIC spectroscopy software (Thermo Scientific) and OriginLab software were used.

The native feedstock was gently extracted from an adult silkworm’s posterior middle silk gland onto a cover glass slide. It was briefly frozen in liquid nitrogen and freeze-dried before measuring it using the FTIR. The silk fibers were degummed similarly to the standard degumming procedure^99^. Briefly, *B. mori* silk cocoons were cut into small pieces and soaked twice, for 15 min each time, in 2 liters of boiling Milli-Q water with 0.02 M sodium carbonate. Next, fibers were exhaustively washed in Milli-Q water and left to dry for three days at room temperature.

## Acknowledgments

Confocal imaging and image analysis were made possible thanks to the de Picciotto Cancer Cell Observatory in Memory of Wolfgang and Ruth Lesser at the Weizmann Institute of Science, and the help of Yoseph Addadi, Ofra Golani, and Tatiana Smirnova. Electron microscopy studies were conducted at the Irving and Cherna Moskowitz Center for Nano and Bio-Nano Imaging at the Weizmann Institute of Science. The authors would like to acknowledge Sharon G Wolf and Nadav Elad for their help with some of the TEM imaging. U.S. acknowledges financial support from, the Nella and Leon Benoziyo Center for Neurological Diseases. In addition, U.S. thanks the Perlman family for funding the Shimanovich Lab at the Weizmann Institute of Science: “This research was made possible in part by the generosity of the Harold Perlman Family.” The authors would like to acknowledge partial support from the Mondry Family Fund for the University of Michigan/Weizmann collaboration, the Gerald Schwartz and Heather Reisman Foundation, and the WIS Sustainability and Energy Research Initiative (SAERI). This research was supported by a research grant from the Tom and Mary Beck Center for Advanced and Intelligent Materials at the Weizmann Institute of Science, Rehovot, Israel. The authors are also grateful to Natalie Page for the English editing.

