## Supplementary data for "The Natural Material Evolution and Stage-wise Assembly of Silk Along the Silk Gland"

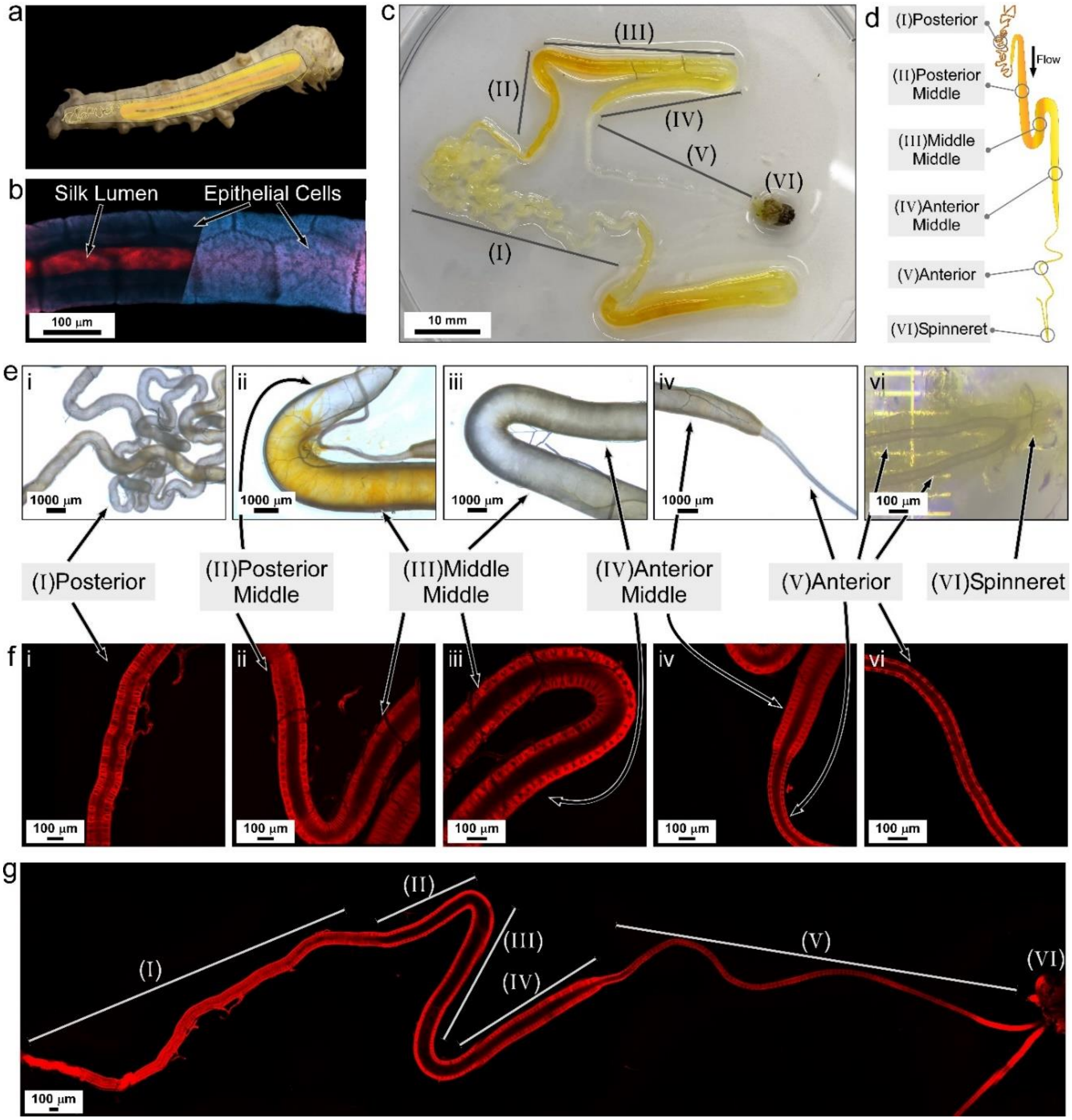


***Figure S1: Silk gland general morphology.*** *(a) B. mori silkworm with illustration of its inside silk gland. (b) The tubular structure of the silk gland (confocal imaging; Nile red staining of anterior section), presenting its constructing epithelial cells and inside lumen. (c) Silk gland of an adult silkworm (5$^{th}$ instra). (d) A drawing of a silk gland indicating its different sections. (e) Selected regions of an adult silk gland (light imaging) . (f) Selected regions and (g) Full length (stitched images) of a nine-day old silk gland (confocal imaging; Nile red staining).*

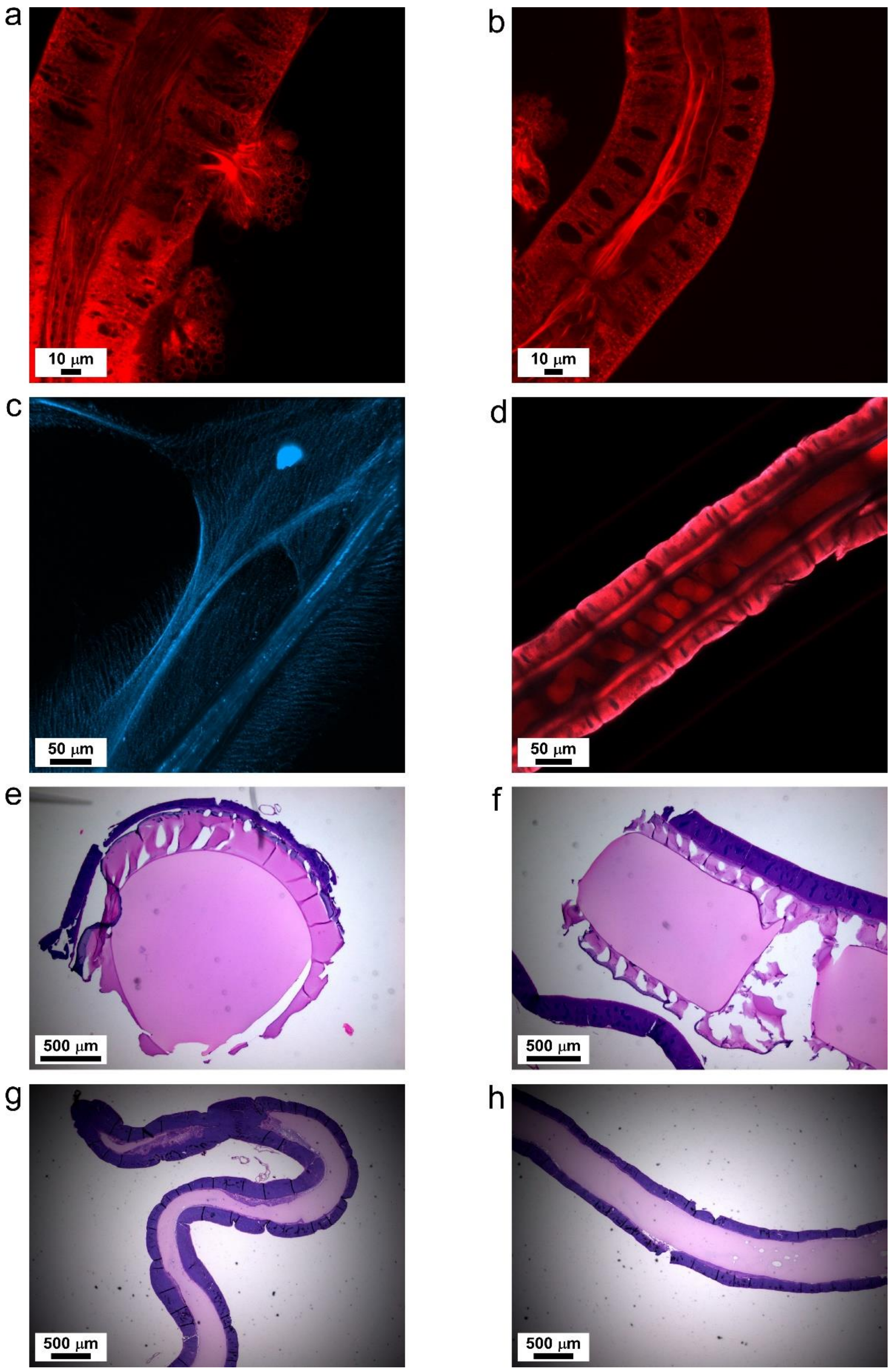


***Figure S2: Distorted silk feedstock.*** *(a,b) Silk feedstock fibrillation inside a nine-days old silkworm silk gland (confocal imaging; Nile red staining), resulted from tissue rupture and unnatural flow. (c) Fibrillation of native silk feedstock, artificially produced due to freeze-drying process (Confocal imaging; interinsic flourescence). Histology of (e,f) middle-middle and (g,h) posterior section of adult silkworm silk gland (H&E staining).*

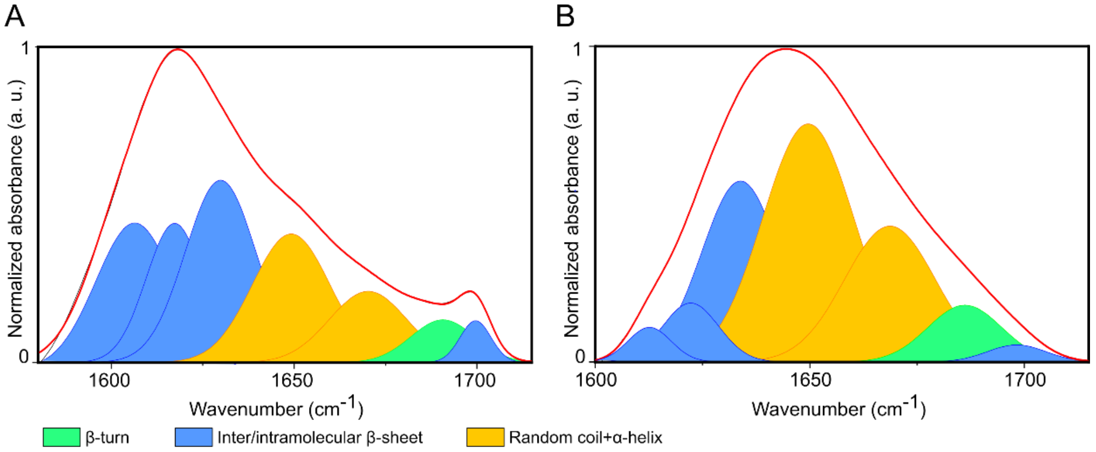


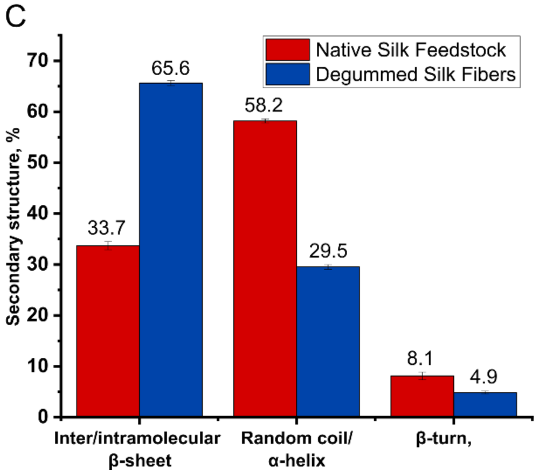


D

| Band position (cm-1) | Assignment |
|---|---|
| 1605 - 1621 | Aggregated strands/β-sheets (weak)[a] |
| 1621 - 1637 | β-sheets (strong)[b] |
| 1638 - 1649 | random coils/extended chains |
| 1650 - 1664 | α−helices |
| 1665 - 1689 | β-turns |
| 1690 - 1705 | β-sheets (weak)[a] |
| [a] Intermolecularβ-sheets [b] Intramolecularβ-sheets | |

***Figure S3:*** *Amide I FTIR spectra of (A) degummed silk fibers and (B) native silk feedstock extracted from the posterior middle section of the silk gland, with a deconvolution analysis depicting the secondary structural composition, including random coils, α-helices, and β-sheets. (C) A comparative analysis of the secondary structure in % of the degummed silk fibers (blue bars) and native silk feedstock (red bars). (D) Table summarizing the FTIR band assignment.*

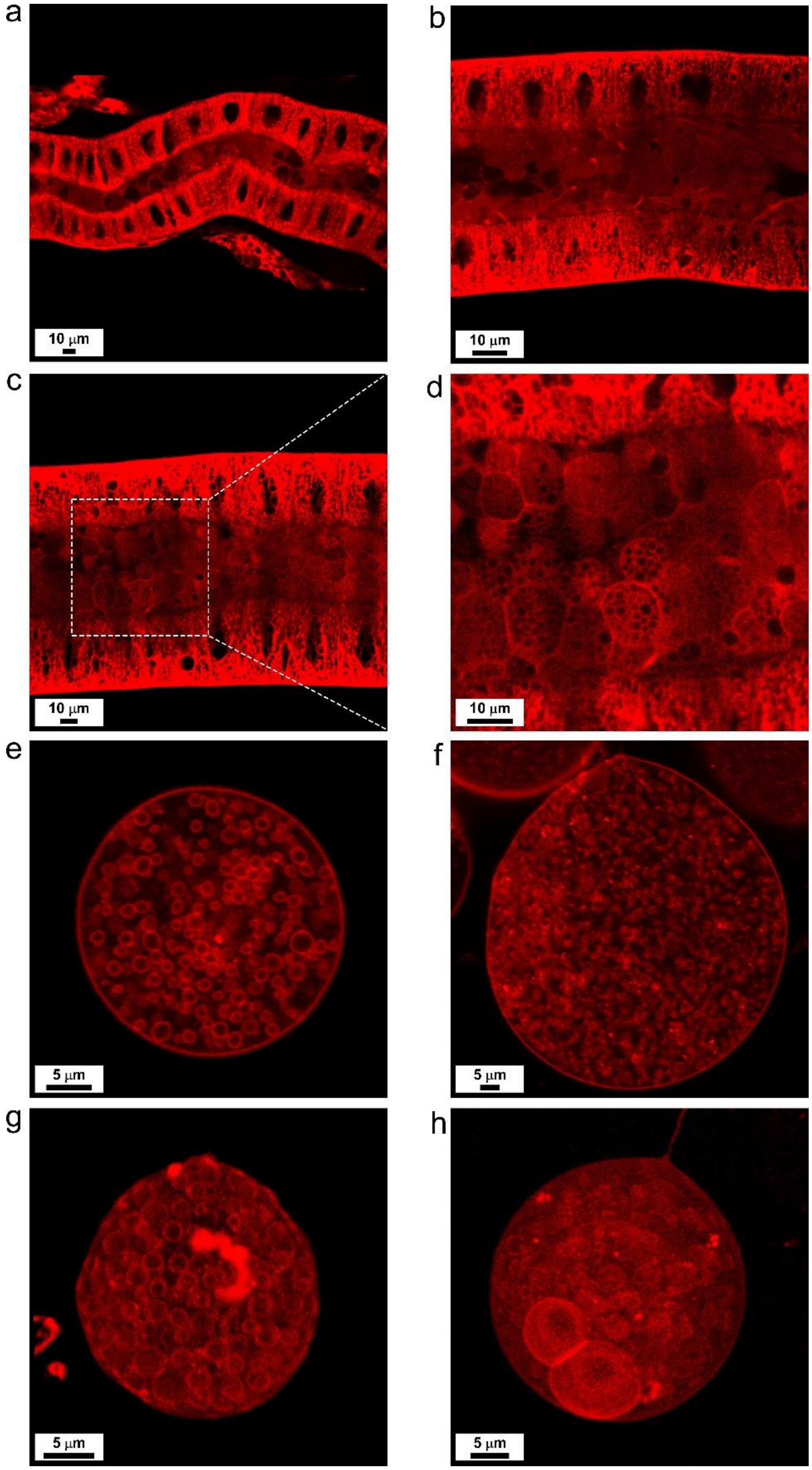


***Figure S4**: **Posterior section's silk feedstock**. (a-d) In-situ confocal imaging of silk feedstock and protein compartments inside the posterior section of the silk gland. (e-i) Ex vivo confocal imaging silk protein compartments from the posterior gland section (Nile red staining).*

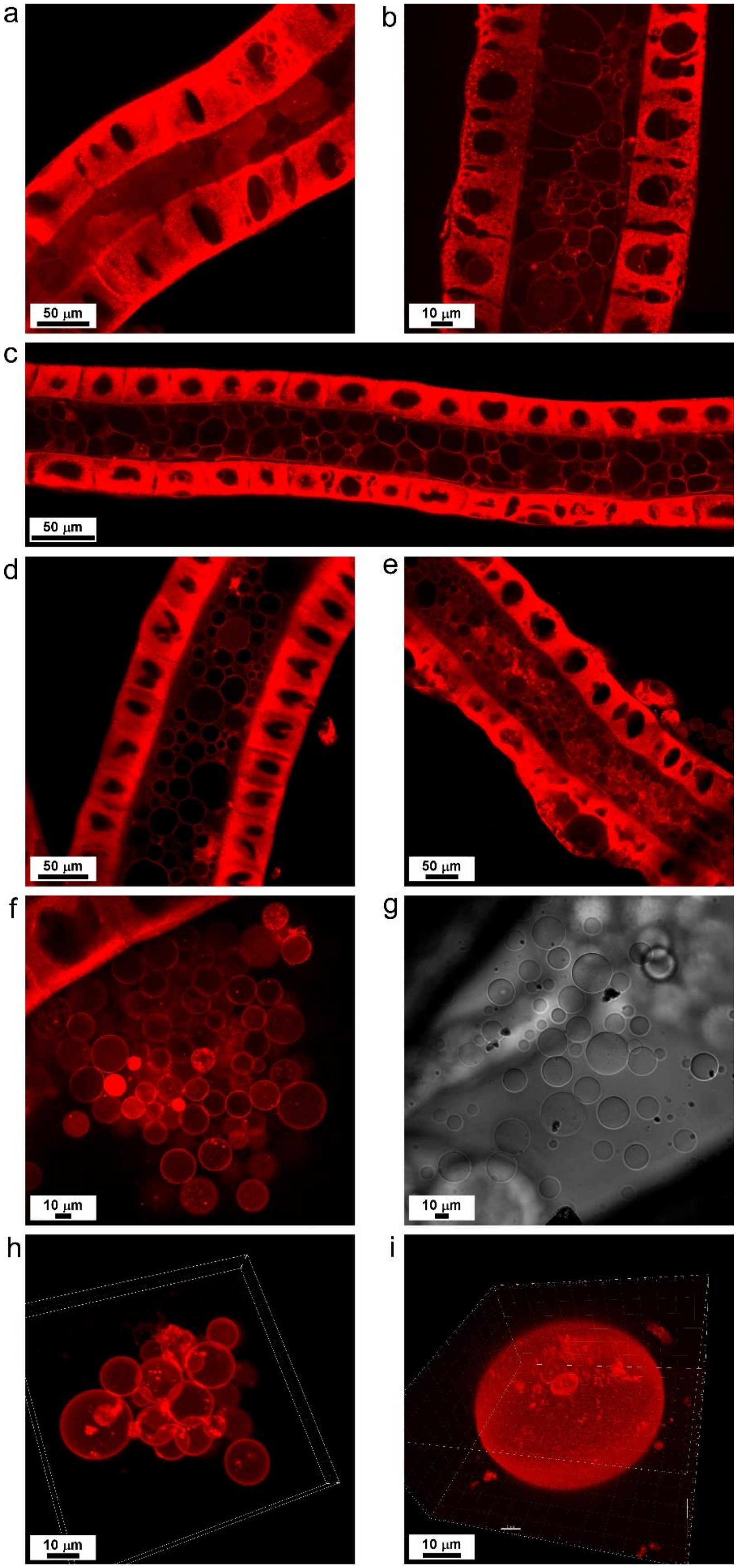


***Figure S5**: **Posterior- and middle-middle sections' of silk feedstock**. (a-d) In-situ confocal imaging of silk feedstock and protein compartments inside the (a-c) posterior-middle and (d,e) middle-middle sections of the silk gland. (f-i) Ex vivo confocal imaging of silk protein compartments from the middle-middle gland section (Nile red staining).*

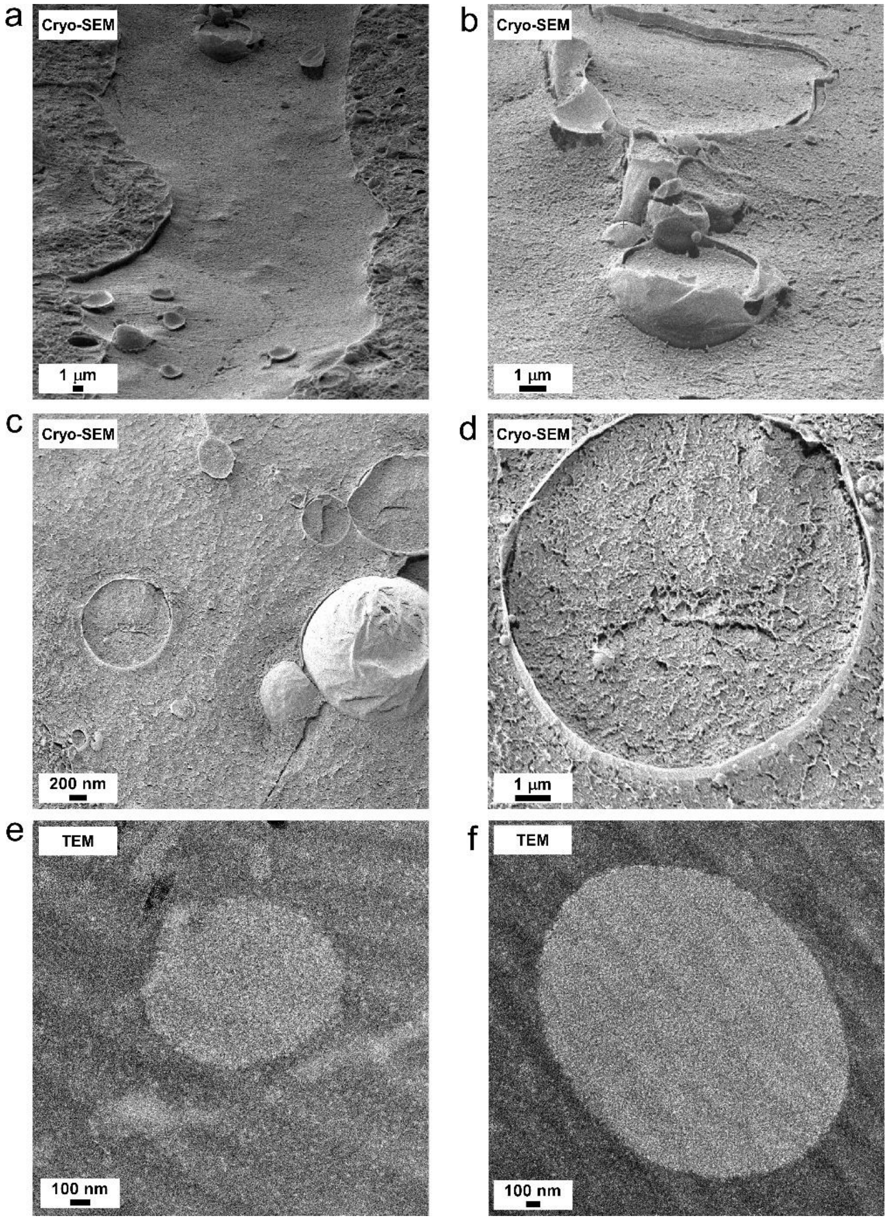


***Figure S6*: *Interior structure of silk protein compartments*.** *(a-d) Cryo-SEM imaging of silk protein compartments from the middle-middle gland section prepared by high-pressure freezing and freeze-fracture. (e,f) TEM images of thin sections from the middle-middle gland section, showing silk protein compartments, prepared by high pressure freezing and freeze substitution.*

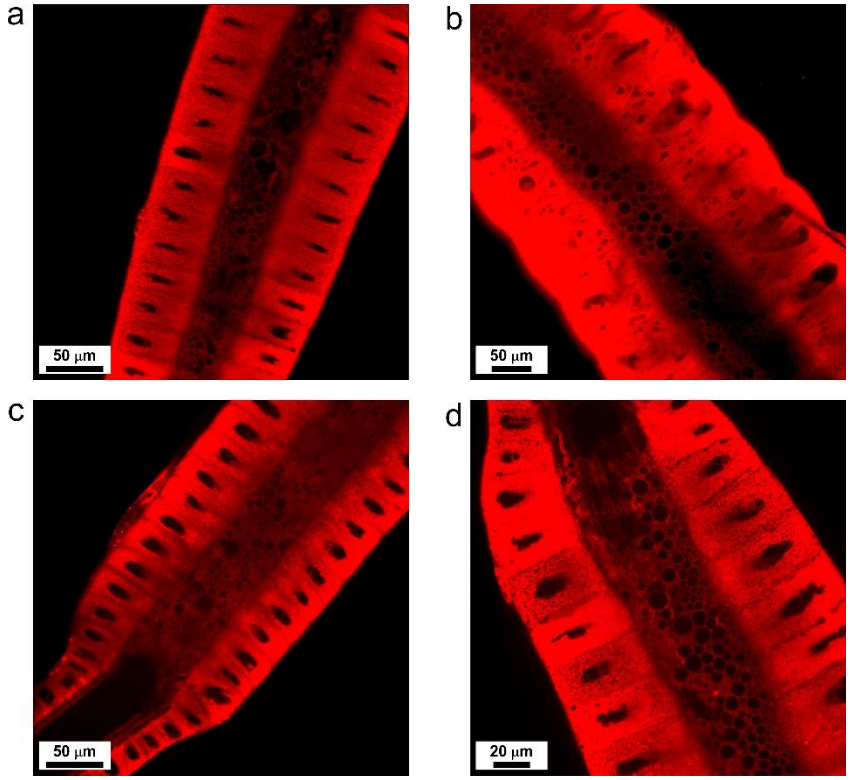


***Figure S7**: **Anterior-middle section silk feedstock**. (a-d) In-situ confocal imaging of silk feedstock and protein compartments inside the anterior-middle section of the silk gland (Nile red staining).*

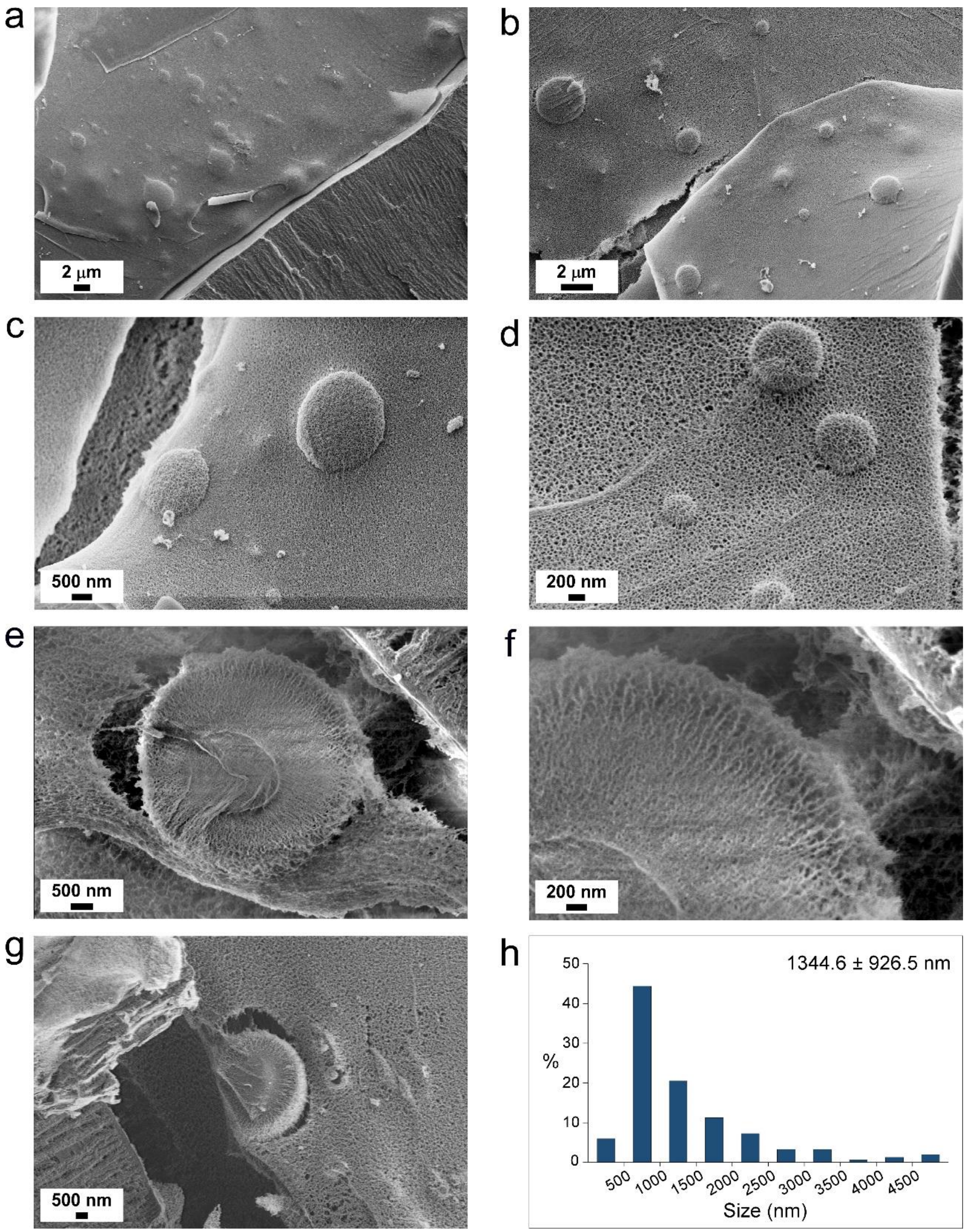


***Figure S8*: *Silk feedstock at the beginning of the anterior gland section*.** *(a-d) In-situ cryo-SEM imaging of silk feedstock at the beginning of the anterior gland section, prepared by high pressure freezing and freeze fracture, shows protein compartments and unstructured feedstock. (e-g) Cryo-SEM imaging of the interior structure of freeze-fractured protein compartments inside the anterior section. (h) Size distribution of protein compartments in the anterior section.*

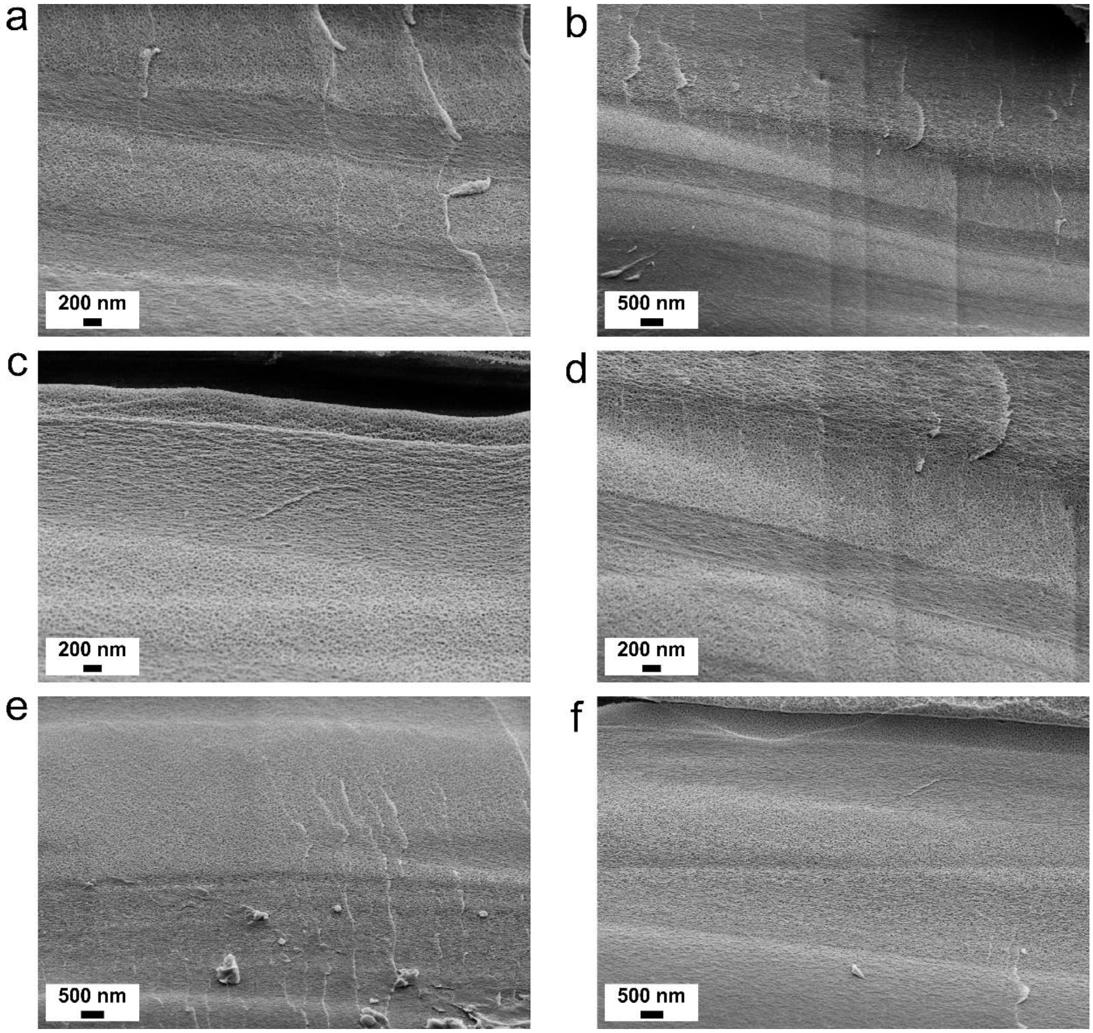


***Figure S9**: **Silk feedstock elongational flow alignment**. (a-e) In-situ cryo-SEM imaging of silk feedstock during molecular alignment inside the anterior silk gland section, prepared by high-pressure freezing and freeze fracture. The imaging shows the protein chain aligning as a response to the elongational flow inside the gland.*

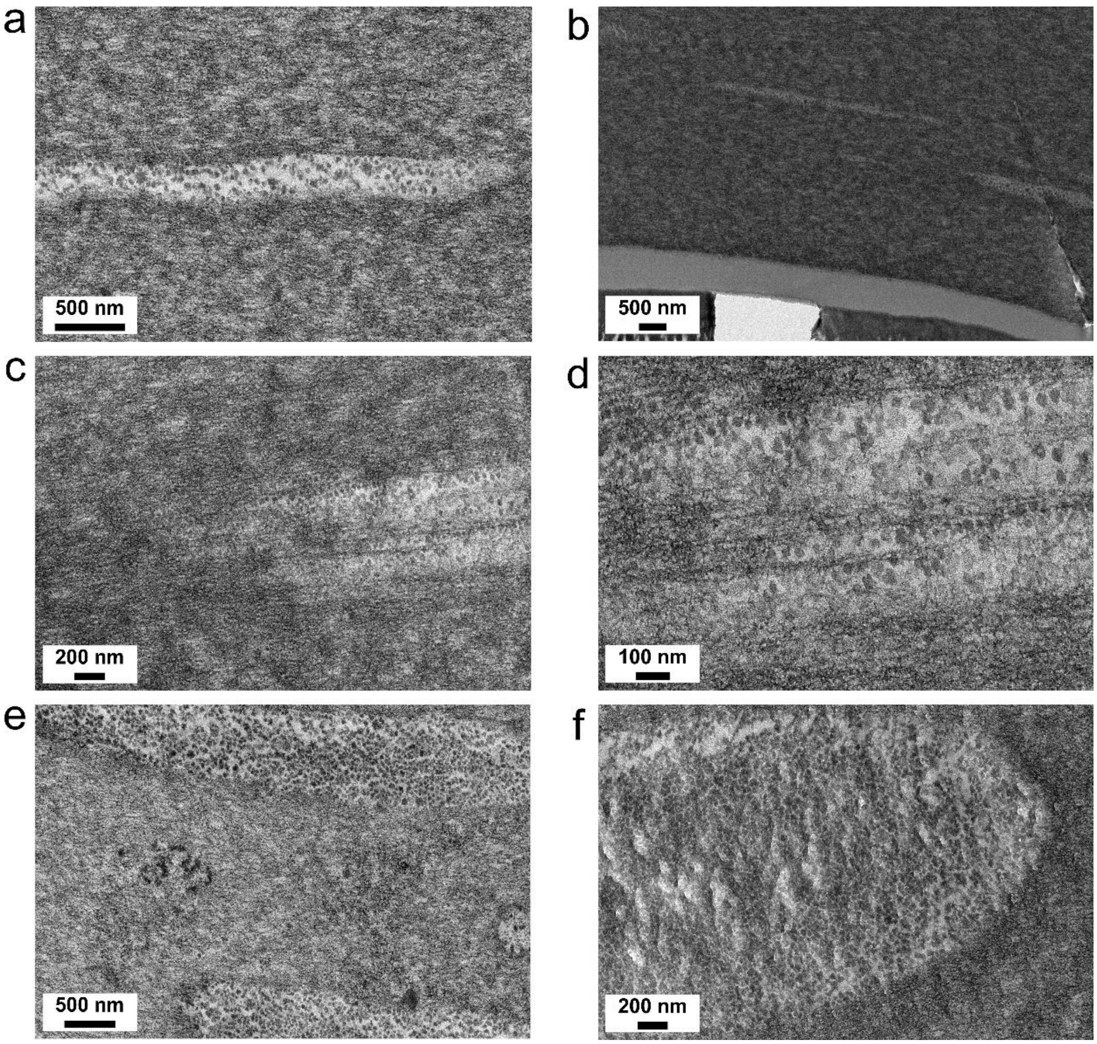


***Figure S10*: *Protein alignment and pahse separation*.** *(a-d) In-situ TEM imaging of silk feedstock during phase separation and alignment inside the anterior silk gland section, , prepared by high-pressure freezing and freeze-subtitution.*

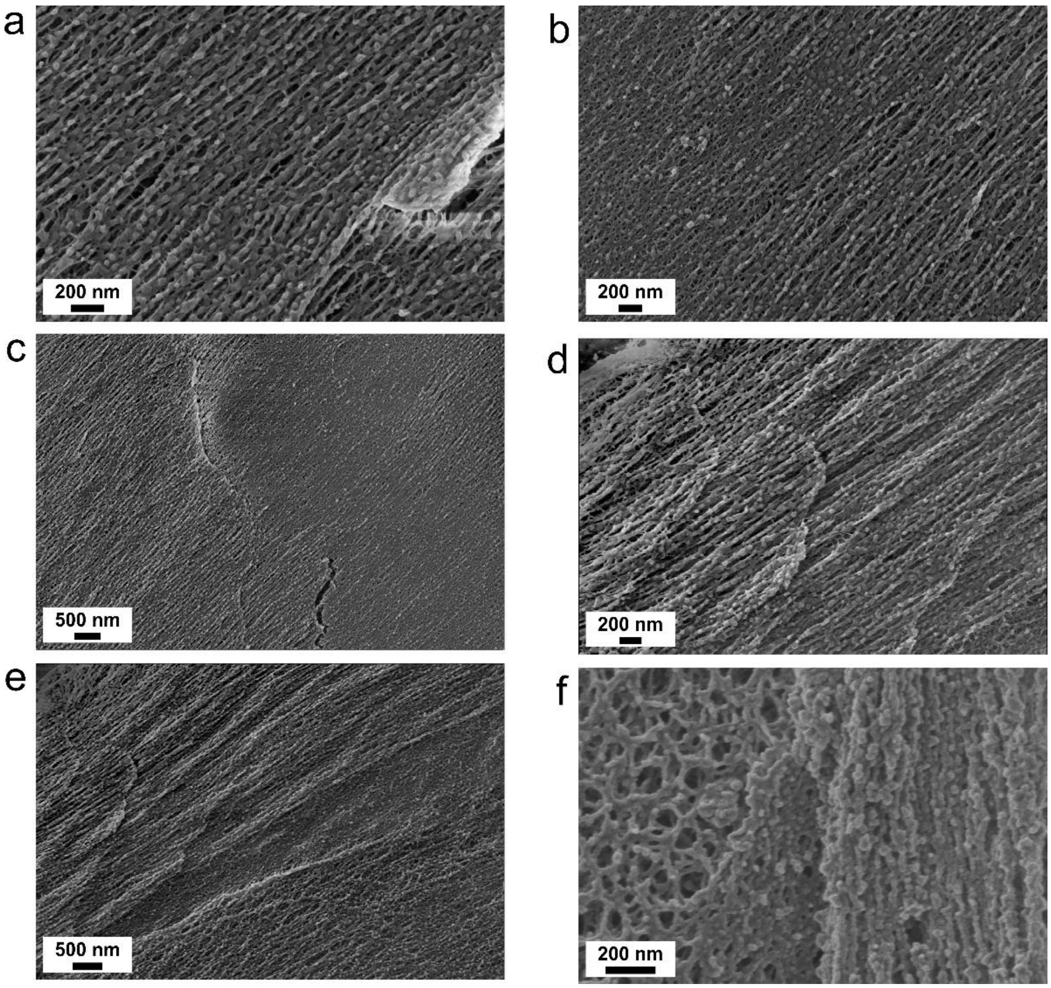


***Figure S11*: *Beads-on-a-string nano-fibrils*.** *(a-f) In-situ cryo-SEM imaging of fibrillated silk feedstock froming a beads-on-a-string shaped nano-fibrils inside the anterior silk gland section, , prepared by high-pressure freezing and freeze fracture.*

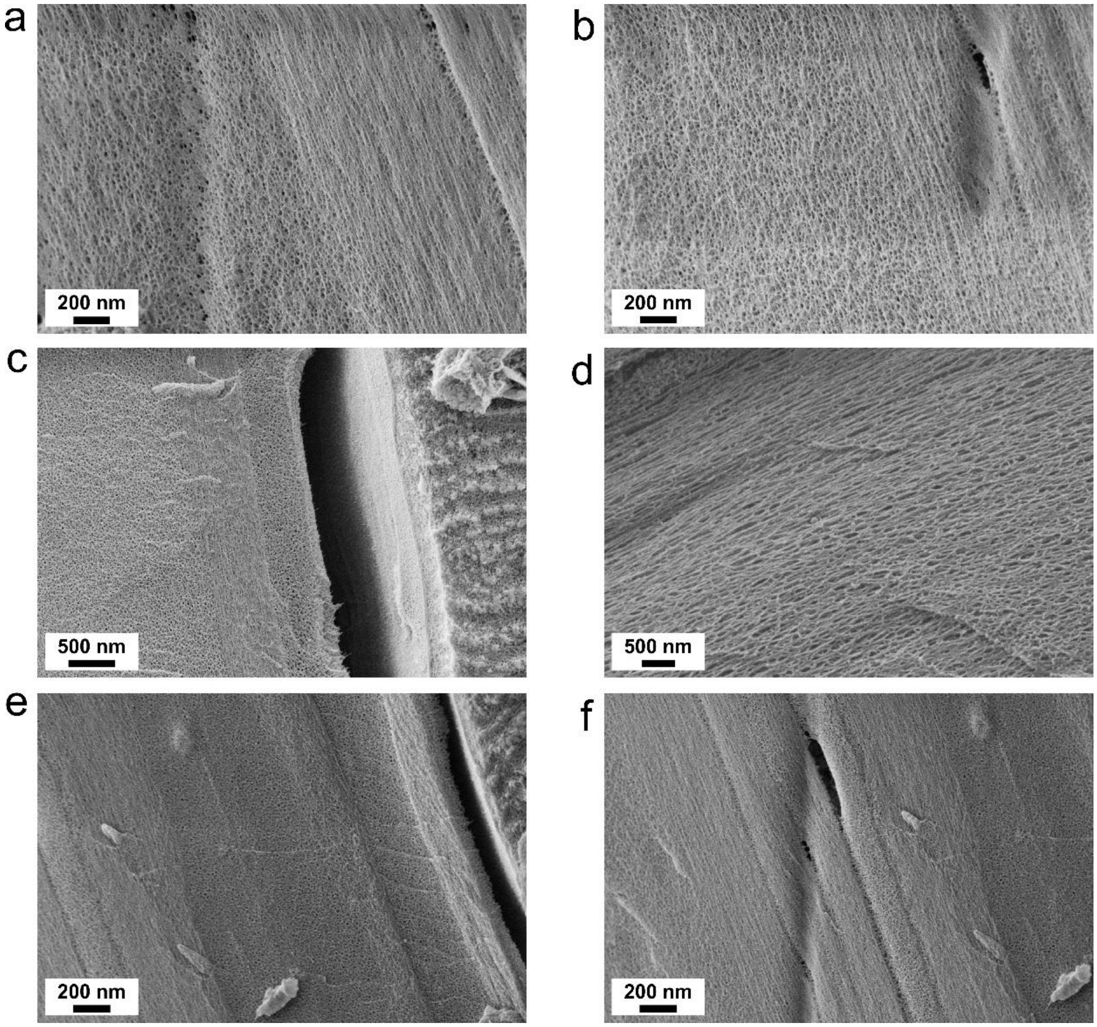


***Figure S12**: **Stretched uniformly-shaped nano-fibrils**. (a-f) In-situ cryo-SEM imaging of silk feedstock fibrillation, forming small, highly-aligned and uniformly-shaped nano-fibrils inside the anterior silk gland section. Samples prepared by high-pressure freezing and freeze fracture.*

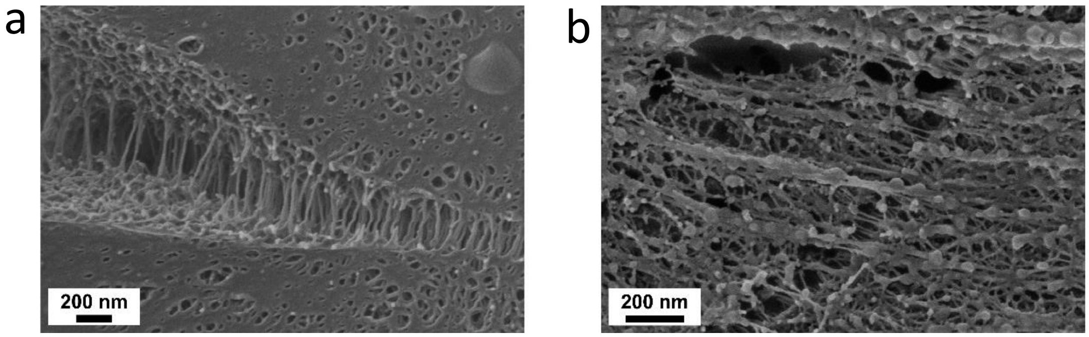


***Figure S13:** (c,d) In-situ cryo-SEM imaging of nano-fibrils stretching due to prolonged sublimation. Samples prepared by high-pressure freezing and freeze fracture.*

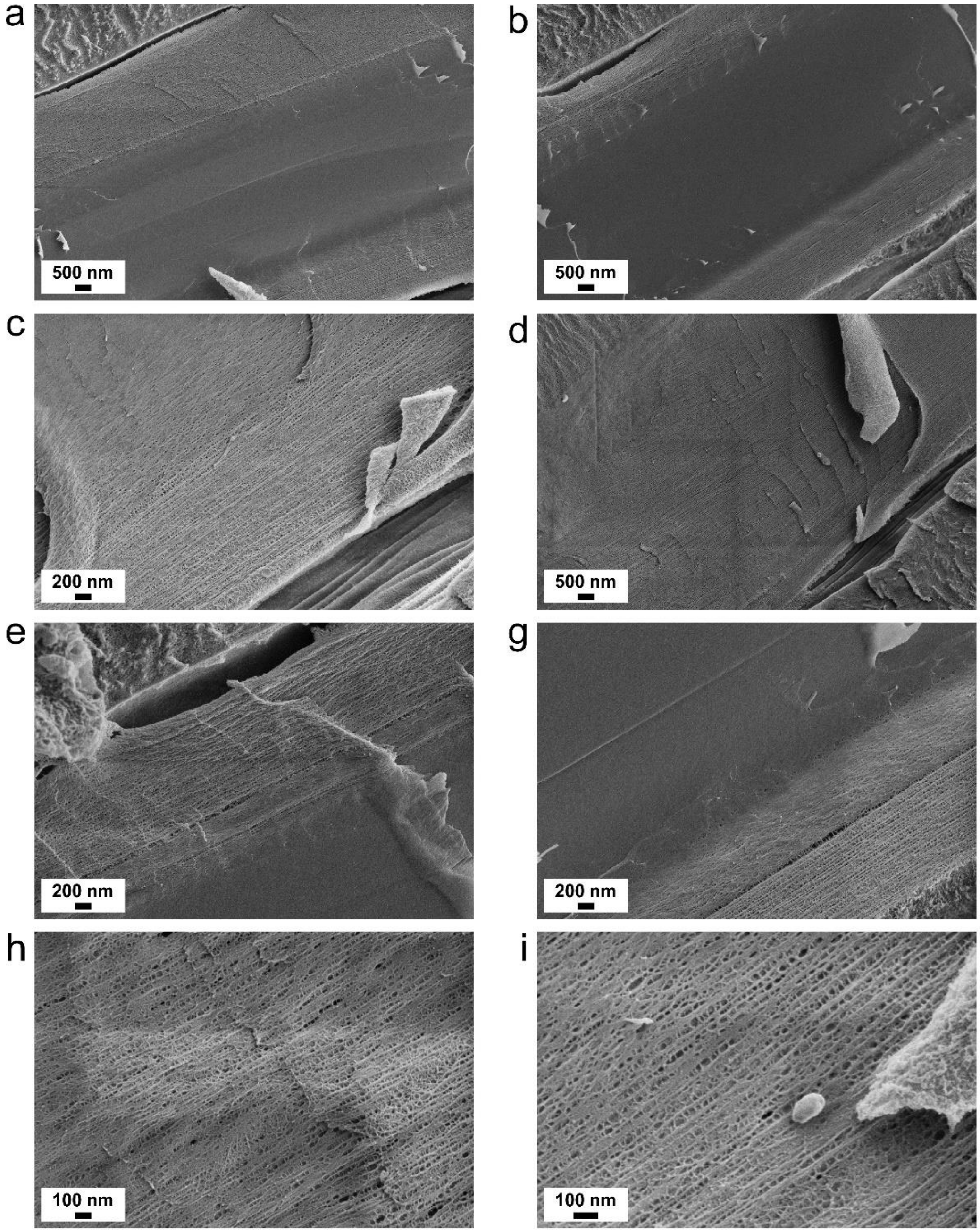


***Figure S14*: *Nano-bundles formation*.** *(a,b) In-situ cryo-SEM imaging of silk feedstock transitioning into nano-bundles inside the front part of anterior silk gland section. (a,b) Overview images of the gland inner tube. (c,d) low magnification images of nano-bundled feedstock. (e,g) Images of the gradual formation process of the nano-bundles. (h,i) High mignification imaging of the nano-bundles morphology. Samples prepared by high-pressure freezing and freeze fracture*

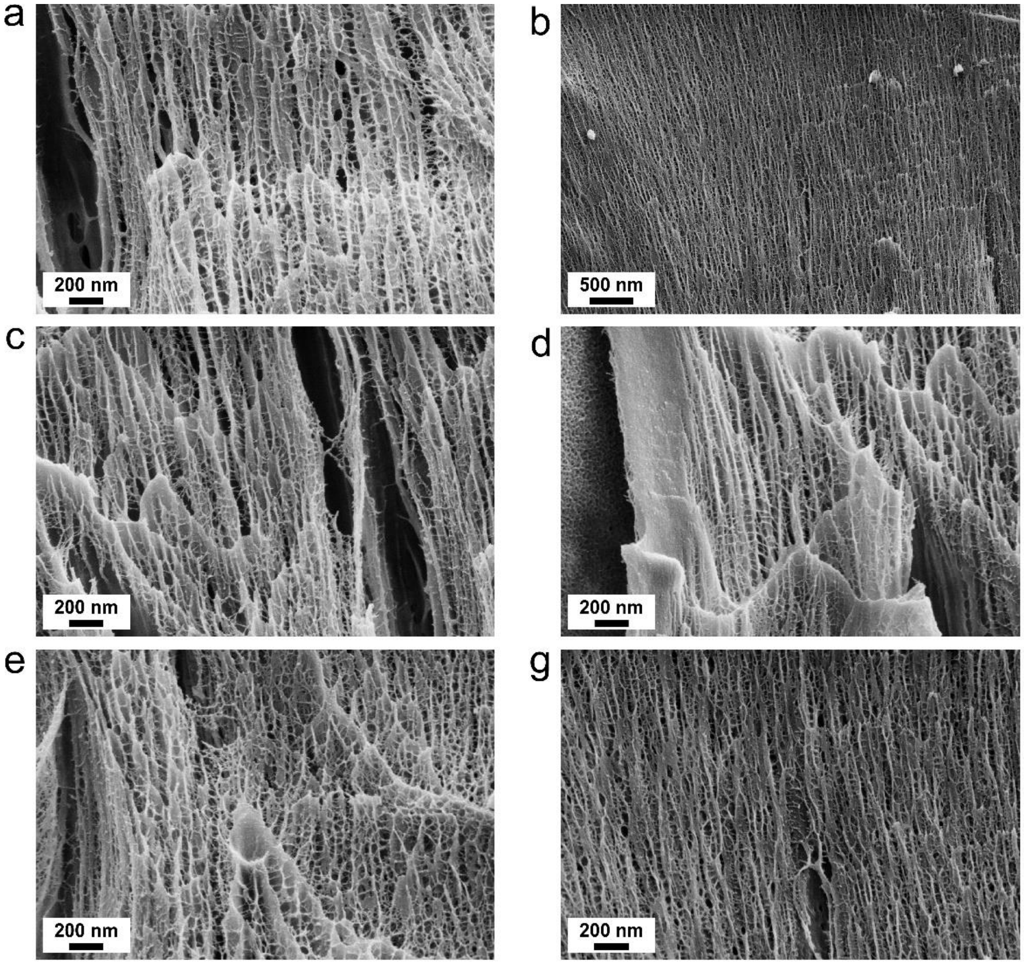


***Figure S15: Further sheared nano-bundles***. *(a-g) In-situ cryo-SEM imaging of silk feedstock nano-bundles after additional high shear flow, prepared by high-pressure freezing and freeze fracture. (h,i) Size distribution of the further sheared nano-bundles and their connecting fibrils. Fibrillated structures diameter distribution analyzed based on their (h) number percent and (i) volume percent.*

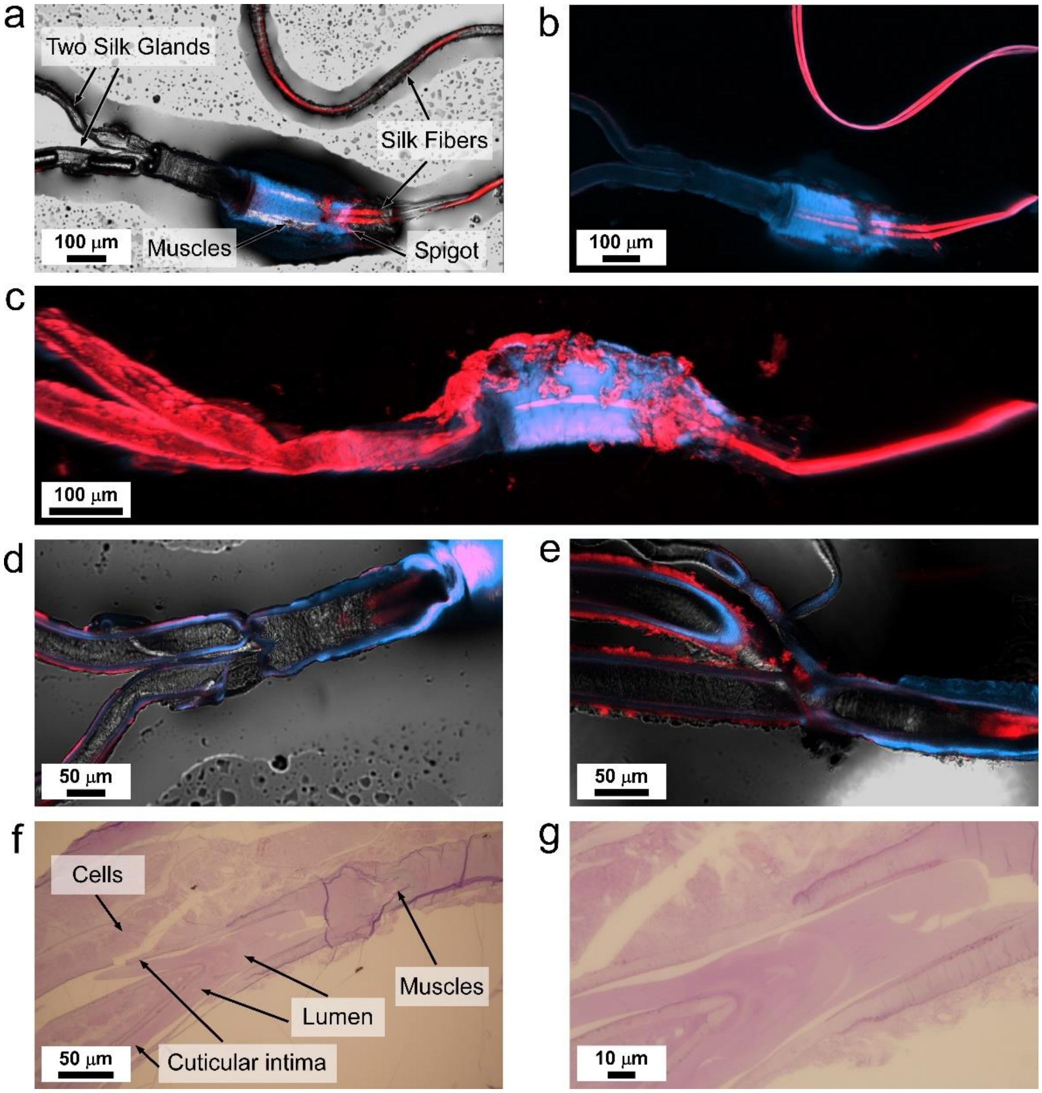


***Figure S16***: *Silk Spinneret. (a-c) Confocal imaging of B. mori spinneret consists of two silk glands merging into one, the silk press muscles (blue), exit spigot, and spun silk fibers (red). Magnified confocal imaging of the two silk glands intersection and merging into one tube before entering the silk press. (f,g) Light imaging of a microtome section of the spinneret and the intersection of the two silk glands, consisting of the epithelial cells, cuticular intima, lumen, and silk press muscles.*

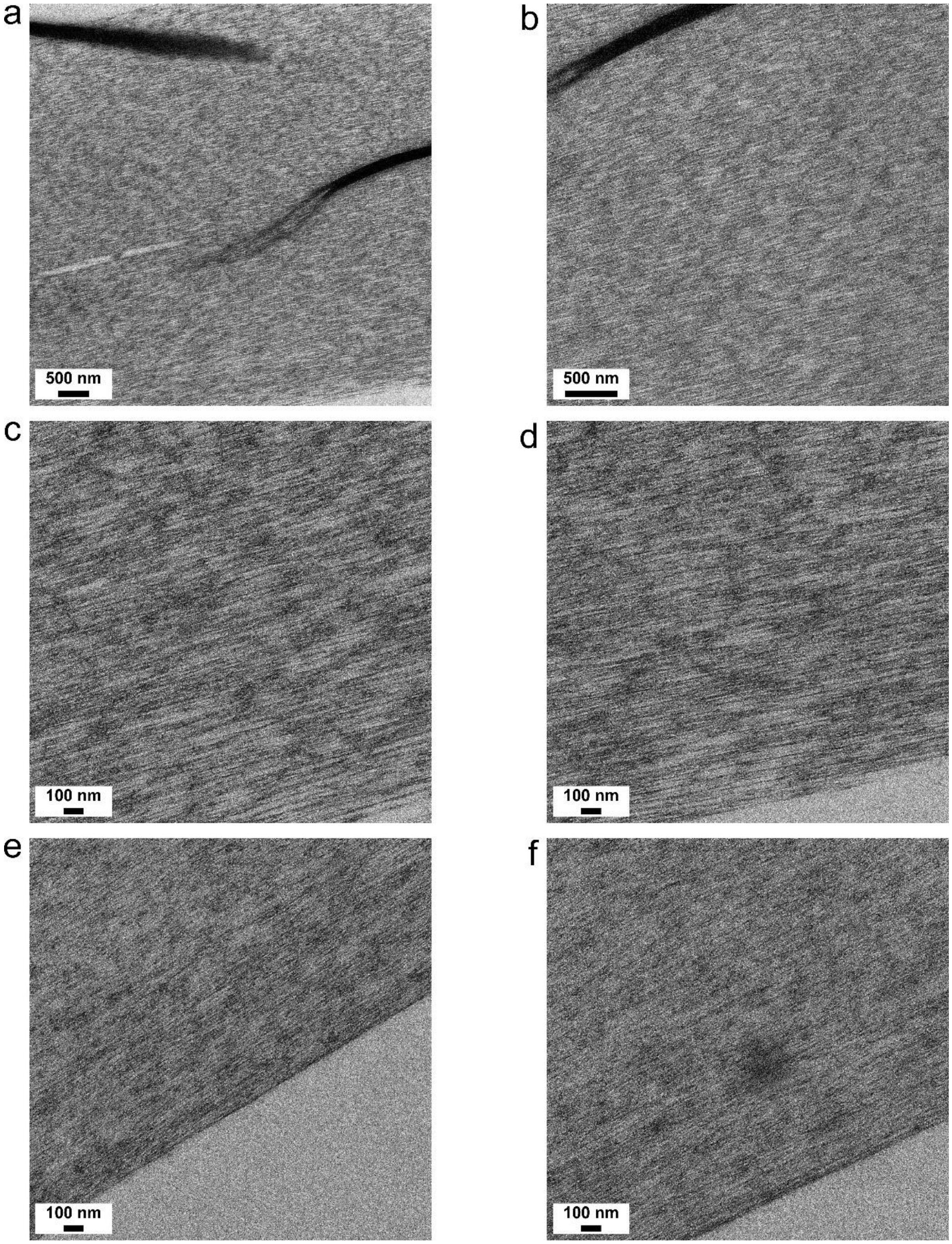


***Figure S17**: **Nano-fibrillated structure of silk feedstock**. (a-f) In-situ TEM imaging of the fibrillated silk feedstock at the front part of the anterior silk gland section, as the two silk glands merge. Samples prepared by high-pressure freezing and freeze-substitution (light images in Figure 20).*

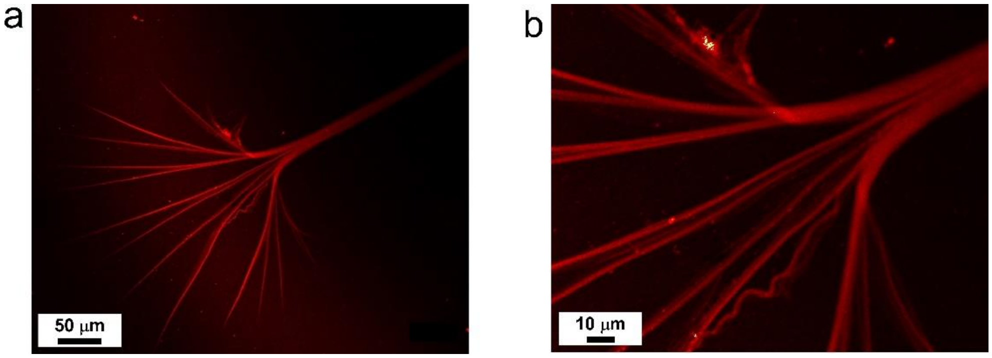


***Figure S18****:(a,b) Confocal imaging of a not fully bundled silk fiber, spun directly from the silk gland in water (Nile red staining).*

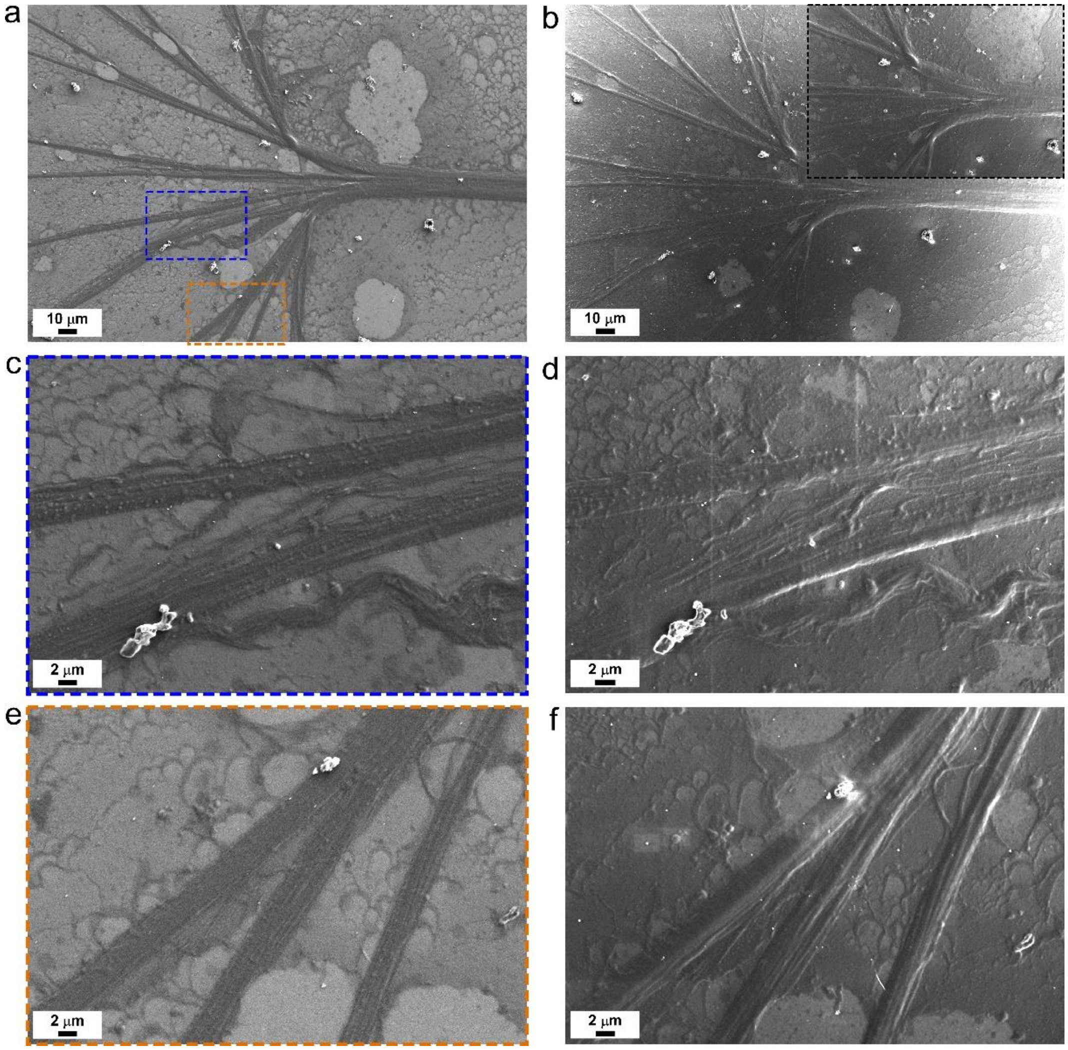


***Figure S19****: SEM imaging of not fully bundled silk fiber. (a,b) overview images and (c-f) magnified images of selected regions of the same fiber were acquired using both (a,c,e) SE2 and (b,d,f) InLens SEM detectors. Inset in (b) is a higher-resolution image with the same scale bar.*

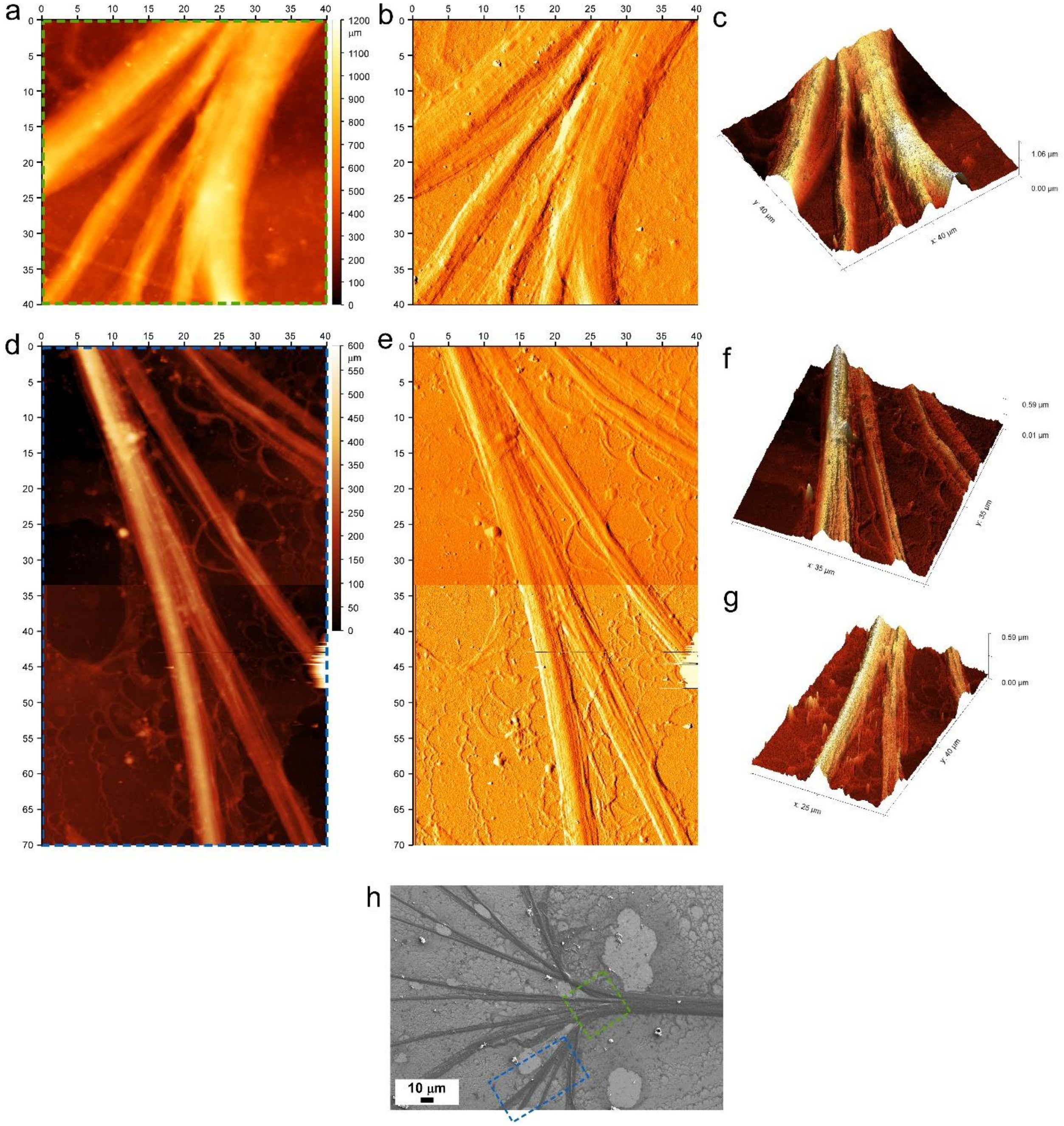


***Figure S20**: **AFM imaging of not fully bundled silk fiber**. AFM (a,d) measured height and (b,e) lock-in amplitude images, and (c,f,g) 3D reconstruction of two scanned regions of the not fully bundled silk fiber. (d) and (e) consists of two stitched images corresponding to the 3D reconstruction in (f) and (g). (h) An overview SEM image of the fiber with marked regions matching the acquired AFM images.*

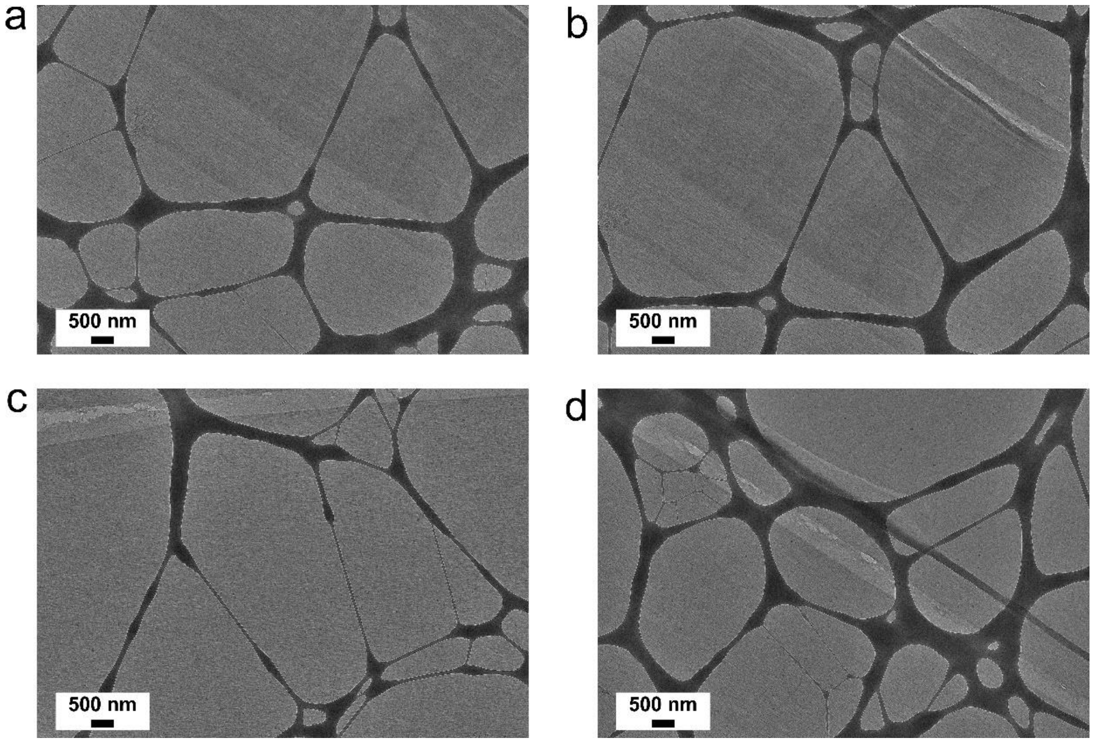


***Figure S21: Nano-structure of native silk fibers***. *(a-d) TEM imaging of the nano-fibrillated structure of native silk fibers, prepared by microtome sectioning of the fibers.*

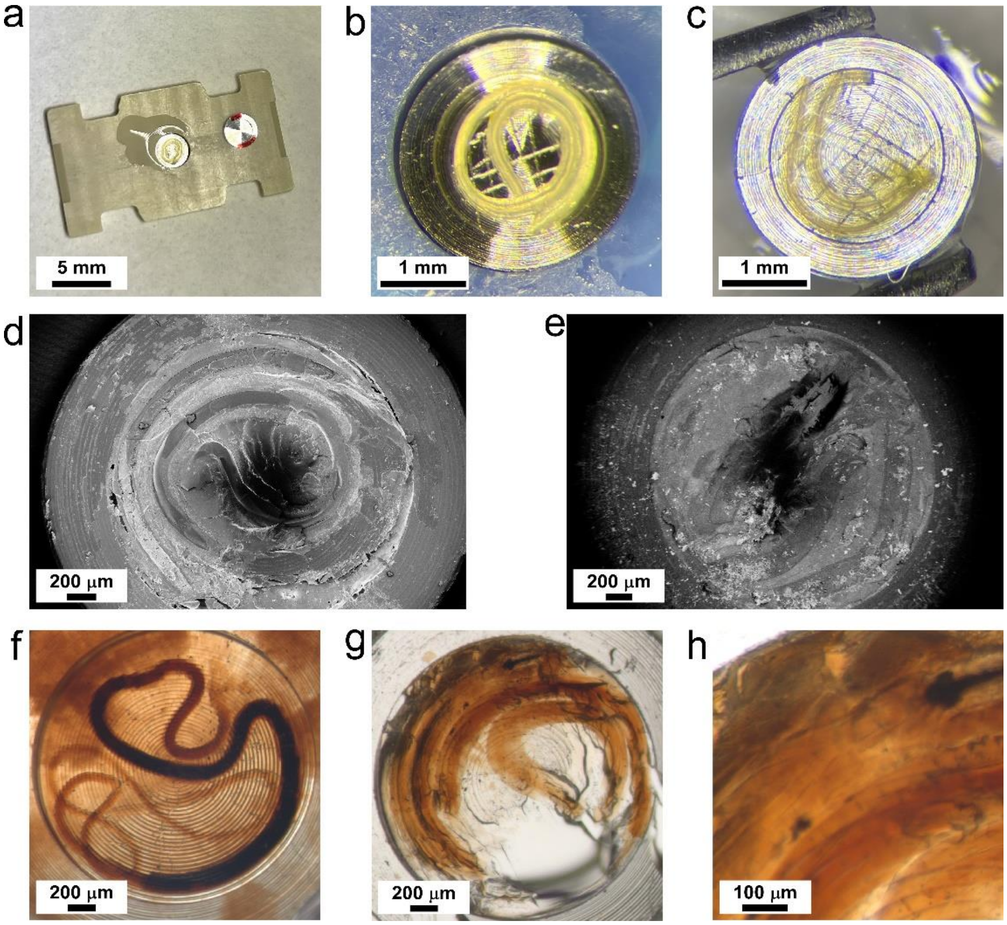


***Figure S22: Cryogenic fixation of silk gland.*** *(a-c) Samples preparation by high pressure freezing (HPF) of adult (5$^{th}$ instra) silkworm silk gland. (d-e) Adult silk glands after HPF and freeze-fracture (Cryo-SEM imaging). Freeze-subtituted samples embeded in epoxy of (f) full length silk gland of a nine-days old silkworm, and (g,h) anterior + spinneret sections of an adult silkworm.*
